# Chromosome-Scale, Telomere-to-Telomere Assembly of Winged Bean (*Psophocarpus tetragonolobus* (L.) DC.) Genome Reveals Lipid Biosynthetic Hotspots and Gene Family Dynamics

**DOI:** 10.1101/2025.08.04.668583

**Authors:** Kishor U. Tribhuvan, Nikhil Kumar Singh, Binay Kumar Singh, Avinash Pandey, Sudhir Kumar, Sujit Kumar Bishi, K. K Kanaka, Tanmaya Kumar Sahu, A. Pattanayak, Vijai Pal Bhadana, Sujay Rakshit

**Author notes:** Present Address: ICAR- Central Institute for Cotton Research, Nagpur (441108), Maharashtra, India. Present Address: ICAR-Indian Institute of Wheat and Barley Research, Karnal (132001), Haryana, India. Authors Contributed equally (Kishor U. Tribhuvan, Nikhil Kumar Singh).

## Abstract

Winged bean (*Psophocarpus tetragonolobus*) is a nutritionally rich but cytologically and genomically underexplored legume with potential for climate-resilient agriculture. We report a telomere-to-telomere, chromosome-scale genome assembly (∼697.69 Mb; N50 = 85.98 Mb; ∼98.25% of the cytometric estimate) of the cultivar AKWB1 using a combination of PacBio HiFi, Illumina, BioNano, and Hi-C technologies. Cytogenetics revealed a diploid karyotype (2n = 18) with exclusively metacentric chromosomes, establishing the first chromosomal ideotype map for the species. The high-quality assembly anchors 98.28% of bases to nine pseudochromosomes and includes 53,745 annotated protein-coding genes, with 97.6% BUSCO completeness. Repeat elements comprise ∼60% of the genome, dominated by LINEs, LTRs, and DNA transposons. KEGG and GO annotations reveal diverse biosynthetic capabilities across metabolic and stress-response pathways. Comparative genomics with nine legumes demonstrates conserved chromosomal synteny with *Glycine max*, absence of recent whole-genome duplication, and a contraction-dominated gene family history. Lipid metabolism analysis identified 750+ genes across 12 KEGG pathways, including fatty acid biosynthesis, elongation, and triacylglycerol assembly. Chromosomes 2, 3, and 5 showed enrichment of lipid biosynthetic loci, suggesting evolutionary hotspots. This reference genome positions *P. tetragonolobus* as a valuable model for studying genome evolution and engineering underutilized legumes for oil yield, protein enrichment, and climate adaptability.

## Main

The winged bean (*Psophocarpus tetragonolobus* (L.) DC.) is a highly nutritious legume that remains largely underused in agriculture and poorly studied at the genomic level, despite its strong potential to contribute to food security in tropical areas and climate-resilient farming^1^. The crop boasts edible seeds, pods, tubers, leaves, and flowers, all of which are rich in protein, oil, and essential amino acids^2^. Its nitrogen-fixing ability and adaptability to marginal tropical environments further highlight its value in sustainable agricultural systems^3^.

Despite nutritional parity with soybean (*Glycine max*) and ecological advantages, winged bean remains absent from mainstream genomic and breeding programs^2^. Prior genome size estimates varied widely, ranging from 580 Mb to over 1.2 Gb^4–6^, reflecting methodological inconsistencies and limiting functional genomics. Furthermore, although a diploid karyotype (2n[=[18) was previously reported^3,7^, a detailed ideogram and chromosome morphology such as the composition of short and long chromosome pairs, remained uncharacterized prior to this study. Advancements in long-read sequencing (PacBio HiFi, Nanopore), chromatin conformation capture (Hi-C), and optical mapping now enable high-contiguity genome assemblies approaching telomere-to-telomere (T2T) resolution in previously recalcitrant plant genomes^8,9^. Leveraging these technologies, we have assembled the chromosome-scale genome of winged bean cultivar AKWB1, anchoring >98% of the estimated genome size. To advance cytogenetic knowledge, we developed an ideotype map and estimated the genome size of the same cultivar.

We further dissected lipid biosynthetic pathways, due to the high seed oil content of the crop, aiming to uncover gene networks governing fatty acid production and remodeling. Comparative genomics with related legumes (*G. max and G. soja*) revealed patterns of genome contraction, conserved synteny, and lineage-specific innovations. This work presents the first cytogenetically validated, reference-grade genome for winged bean, establishing a platform for trait discovery, molecular breeding, and orphan legume improvement.

## Results

### Genome size estimation and chromosome karyotyping

The genome size of *P. tetragonolobus* cv. AKWB1 was estimated to be approximately 710.94 Mb through flow cytometric analysis, employing *Oryza sativa* (430 Mb) as an internal reference standard (Supplementary Fig. S1). Karyotype analysis revealed nine homologous chromosome pairs exhibiting uniform morphology (Fig. 1a-b; Supplementary Table S1). Based on arm ratio (p/q), all chromosomes were classified as metacentric, with values ranging from 1.03 to 1.79^10^. Chromosome 1 exhibited the greatest inter-homolog length variation, with lengths ranging from 182.08 µm to 200.74 µm (mean ± s.e.: 191.41[±[6.60[µm). Detailed morphometric measurements for all chromosomes are provided in Supplementary Table S1.

**Figure 1.**
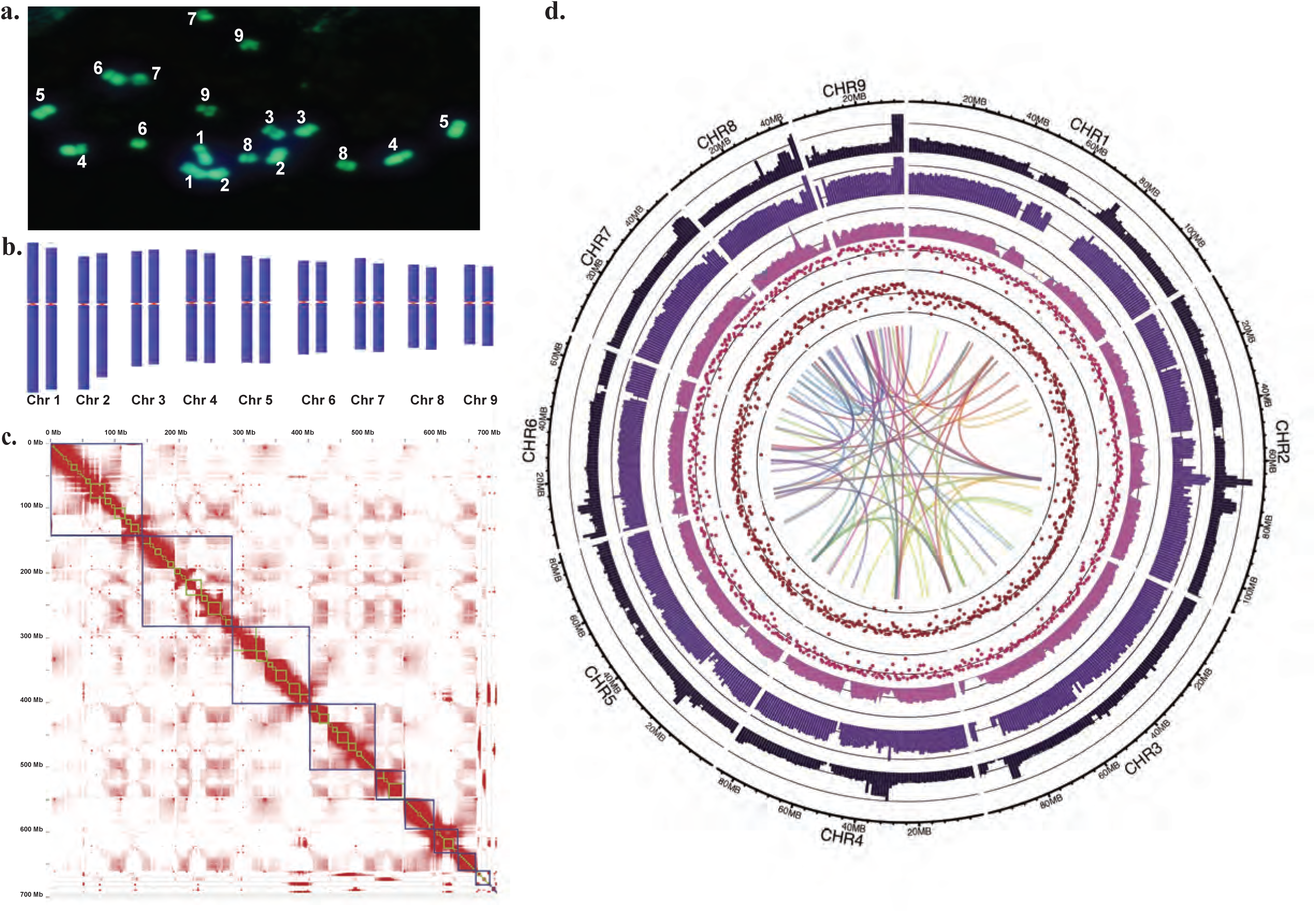
Chromosomal architecture and genome-wide organization of the *P. tetragonolobus*. (a) Fluorescence microscopy image of a metaphase chromosome spread showing 18 chromosomes (9 homologous pairs), labelled 1–9 (b) Ideotype representation of the nine chromosome pairs, arranged by decreasing size and annotated with centromere positions to illustrate overall chromosome morphology (c) Hi-C contact matrix displaying genome-wide chromatin interaction frequencies. Interaction intensity is represented by shades of red, with darker regions indicating higher contact frequency. Blue boxes demarcate individual chromosomes and topologically associating domains (TADs) (d) Circos plot summarizing genome-wide structural and compositional features of P. tetragonolobus. From outermost to innermost, the tracks show: (1) chromosome scale bars proportional to physical length;(2) GC content (black, 1 Mb bins);(3) gene density (purple);(4) total transposable element (TE) density (dark violet);(5) LINE element density (pink); and (6) LTR element density (red).The central colored ribbons connect syntenic regions across chromosomes, revealing conserved intra-genomic rearrangements. This composite view highlights regional variation in GC content, gene/repeat distribution, and evolutionary collinearity.

### Chromosome-scale genome assembly

Chromosome-scale, high-quality genome assembly of *P. tetragonolobus* was achieved by integrating PacBio HiFi long reads, Illumina short reads, BioNano optical maps, and Hi-C chromatin conformation capture data. We obtained ∼990.68[Gb of HiFi reads (∼1,415× coverage; mean subread length: 182,250 bp), 311.35[Gb of Illumina reads (338× coverage; mean read length: 150 bp), 45.3[Gb of Hi-C data (65× coverage), and ∼652.3[Gb of BioNano data (N50: 184.9[Kb) (Supplementary Table S2). De novo assembly using PacBio HiFi reads yielded a primary assembly of 862.2[Mb in 1,294 contigs (N50: 14.62[Mb; L50: 18) with a BUSCO completeness of 97.2% and 8.5% duplicated orthologs, likely due to haplotypic redundancy (Supplementary Table S3). After purging, the refined assembly was reduced to 716.85[Mb across 165 contigs (N50: 16.93[Mb), and duplication dropped to 4.0%. Illumina reads were used for error correction, improving base-level accuracy. Integration of BioNano maps using the Hybrid Scaffold pipeline improved assembly continuity to 59 scaffolds (N50: 19.36[Mb; max: 88.79[Mb), with 97.0% BUSCO completeness (Supplementary Table S3). Hi-C scaffolding anchored the final assembly into 15 scaffolds totaling 697.69[Mb (98.25% of the estimated genome), of which 685.69[Mb (98.28% of the assembly) was assigned to nine pseudochromosomes corresponding to the haploid karyotype (Table 1, Fig. 1c), and the remaining 14.8[Mb was distributed across six unplaced scaffolds. The pseudochromosomes ranged in size from 34.86[Mb to 111.65[Mb (Supplementary Table S4). Telomeric repeat motifs (TTTAGGG)n were detected at one or both ends of all nine pseudochromosomes (Supplementary Fig. S3), indicating near-complete chromosomal assemblies. The final assembly exhibited high contiguity (N50: 86.0[Mb; L50: 4; N90: 41.8[Mb; L90: 8), low gap density (111 gaps; 27.5[Ns/100 kb), and a GC content of 31.08%. BUSCO analysis revealed 97.61% completeness, including 94.5% single-copy genes, highlighting the assembly’s completeness and structural accuracy (Table 1). The genome assembly and annotations have been deposited in the NCBI Genome database and are publicly available under BioProject PRJNA1169650 accession number JBMOEB000000000.

### Annotation of the genome

Repeat annotation revealed that 60% of the genome (409.66[Mb) comprises repetitive elements (Fig. 1d; Supplementary Table S5), with LINEs (32.33%), LTRs (11.3%), and DNA transposons (9.31%) being the most abundant. Unclassified repeats (7.06%) may represent novel families. Small RNA-associated repeats, SSRs, and low-complexity regions contributed 5.66%, 4.2%, and 2.4%, respectively, while satellite repeats were rare (<0.01%) (Fig 1d; Supplementary Table S5). Comprehensive gene prediction identified 53,745 protein-coding genes, including 49,431 anchored to the nine pseudochromosomes (Fig. 1d; Table[1; Supplementary Table S6). Gene lengths ranged from 87 bp to 358,828 bp (mean: 2,684.91[bp; median: 1,444 bp), with an average of 4.72 exons per gene and a mean exon length of 347.1[bp. Coding sequences (CDSs) had a mean length of 1,148.38 bp and ranged from 24 bp to 347,019 bp, with 62.97% being shorter than 1,000[bp (Supplementary Table S5; Supplementary Fig. S4).

A multi-layer annotation revealed that 74.3% of genes showed homology to National Center for Biotechnology Information Non-Redundant (NCBI NR) database, 46.9% contained Pfam domains, and 60.0% were assigned Gene ontology (GO) terms (Fig. 2a; Supplementary Table S7). Evolutionary genealogy of genes: Non-supervised Orthologous Groups (EggNOG)-based orthology analysis annotated 70% of genes, with 31.28% assigned GO terms, 13.1% linked to Enzyme Commission (EC) numbers, and 28.5% mapped to KEGG Orthology (KO) pathways (Supplementary Table S7). GO annotation assigned 49,566 Biological Process, 42,030 Molecular Function, and 31,936 Cellular Component terms across over 21,000 genes, highlighting the functional diversity of the genome (Fig. 2b–d). Enzyme classification based on EC numbers revealed transferases as the most abundant class (7,299 genes), followed by hydrolases (5,986) and oxidoreductases (2,858), reflecting broad biosynthetic and degradative potential (Fig. 3a). Of the 49,431 genes on the nine core chromosomes, 10,217 were annotated with KO identifiers via EggNOG-mapper, enabling classification into 127 KEGG pathways, including secondary metabolism, amino acid metabolism, hormone signaling, and stress responses. The most gene-enriched pathways included ‘Metabolic pathways’ (ko01100; 2,998 genes), ‘Secondary metabolite biosynthesis’ (ko01110; 1,759), Biosynthesis of lipids (ko1004; 1130 genes), Starch and sucrose metabolism (ko00500; 485 genes) (Fig. 3b; Supplementary Table S8).

**Figure 2.**
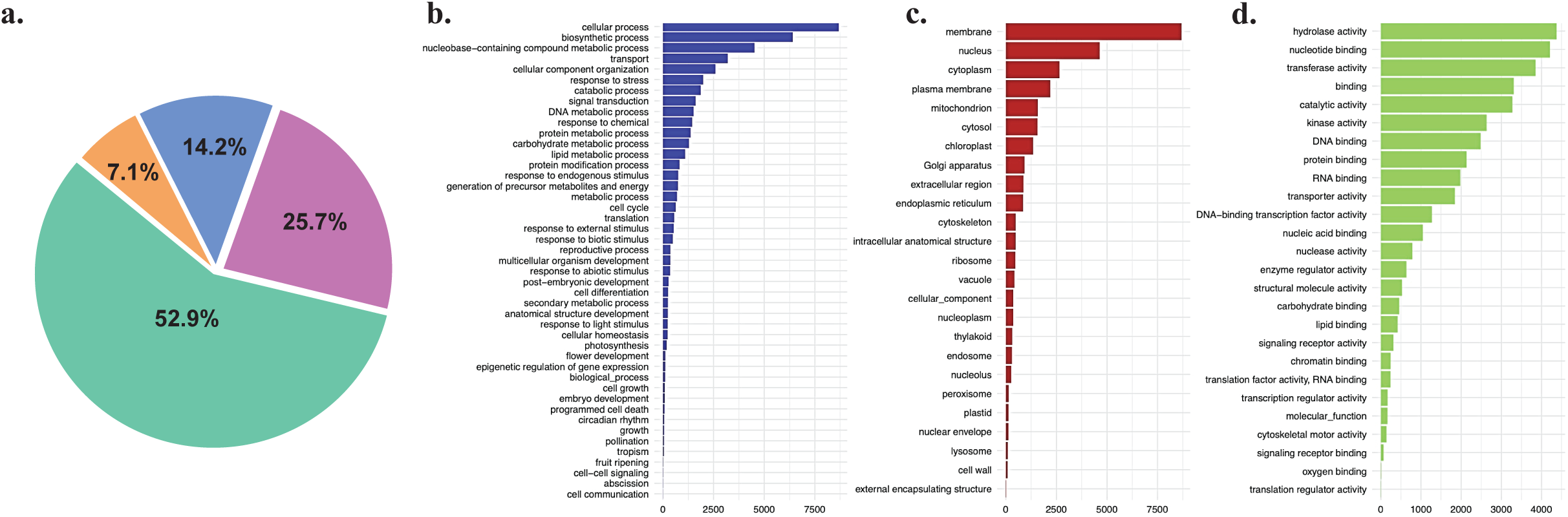
Functional annotation and enrichment of enzymes in *P. tetragonolobus*. (a) Pie chart summarizing the functional annotation status of predicted genes in P. tetragonolobus. A total of 52.9% of genes show significant homology to known proteins based on BLAST searches, 25.7% remain unannotated with no BLAST hits, 14.2% are annotated with Gene Ontology (GO) terms, and 7.1% possess both GO and GO-Slim annotations, indicating a subset of genes with higher-level functional classification. (b) GO terms under the Biological Process (BP) category indicate a high representation of genes involved in metabolic and biosynthetic processes, signal transduction, responses to various stimuli (including biotic, abiotic, and chemical), developmental processes, and cellular organization. (c) GO terms in the Cellular Component (CC) category reveal widespread subcellular localization of enzymes, predominantly in the nucleus, cytoplasm, plasma membrane, mitochondria, chloroplasts, and other intracellular structures. (d) Molecular Function (MF) annotations highlight dominant roles such as catalytic activity, nucleotide binding, hydrolase and transferase activity, and protein binding.

**Figure 3.**
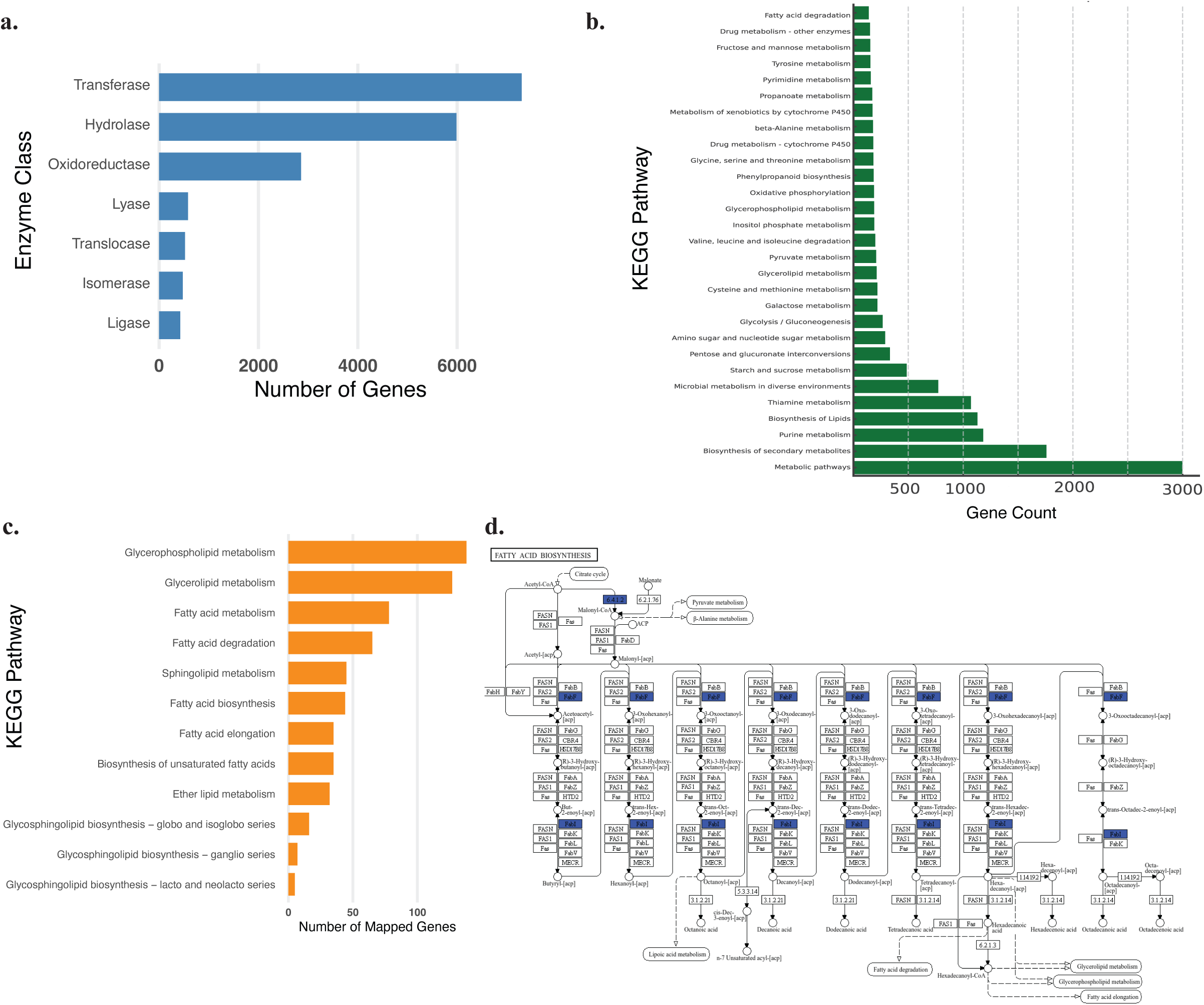
Functional classification of annotated genes with emphasis on lipid metabolism in *P. tetragonolobus*. (a) Enzyme class distribution of annotated genes. Functional classification of genes into the seven major EC (Enzyme Commission) classes based on KEGG and InterProScan annotations. The plot highlights the dominance of transferases, hydrolases, and oxidoreductases, reflecting the genome’s broad enzymatic potential and metabolic flexibility. (b) Bar plot showing the number of genes assigned to major KEGG metabolic pathways based on orthology mapping using EggNOG-mapper. A total of 30 pathways are shown, sorted in descending order of gene count, and labeled with their respective KEGG identifiers (koXXXX). The most highly represented pathway was Metabolic pathways (ko01100) with 2,998 genes, followed by Biosynthesis of secondary metabolites (ko01110; 1,759 genes), and Biosynthesis of lipids (ko00061; 1,130 genes), reflecting the species’ broad biosynthetic potential. Lipid-related pathways such as Glycerolipid metabolism (ko00561; 212 genes), Glycerophospholipid metabolism (ko00564; 189 genes), and Fatty acid degradation (ko00071; 140 genes) are prominently represented, highlighting the metabolic investment in oil biosynthesis and remodeling. Several carbohydrate metabolism pathways—including Glycolysis/Gluconeogenesis (ko00010), Starch and sucrose metabolism (ko00500), and Pentose and glucuronate interconversions (ko00040)—are also well populated, underscoring the active primary metabolic framework. This functional annotation provides insight into the genome-wide distribution of enzymatic potential in P. tetragonolobus, supporting its utility for trait dissection and metabolic engineering. (c) Mapping of genes to lipid and fatty acid metabolism pathways. This panel presents gene counts associated with lipid-related KEGG pathways including fatty acid biosynthesis (ko00061), elongation (ko00062), degradation (ko00071), glycerolipid metabolism (ko00561), glycerophospholipid metabolism (ko00564), and biosynthesis of unsaturated fatty acids (ko01040). Gene assignment reveals robust representation across all major lipid biosynthetic and remodeling routes, supporting the species’ high seed oil content. (d) Gene involvement in sphingolipid and glycosphingolipid pathways. Specific pathways involved in complex lipid biosynthesis are shown, including sphingolipid metabolism and three subclasses of glycosphingolipid biosynthesis (lacto/neolacto series, globo/isoglobo series, and ganglio series). These highlight the structural lipid diversity encoded in the P. tetragonolobus genome, potentially relevant for membrane stability and signaling.

### Genome-wide KEGG pathway mapping reveals candidate genes for oil biosynthesis

Given its phylogenetic proximity to *G. max* and high seed lipid content, *P. tetragonolobus* was analyzed for its oil biosynthesis potential through genome-scale KEGG pathway profiling. A total of 12 lipid-related KEGG pathways were identified, encompassing over 750 genes (Figure[3c; Supplementary Tables S8–S9). Among these, glycerophospholipid metabolism (ko00564) was the most gene-enriched pathway with 138 genes, followed by glycerolipid metabolism (ko00561) with 127 genes. Fatty acid metabolism (ko00071; 78 genes) and fatty acid degradation (ko00071; 65 genes) were also detected, indicating the presence of both biosynthetic and catabolic modules. Additional biosynthetic routes, including Sphingolipid metabolism (ko00600; 47 genes), fatty acid biosynthesis (ko00061; 44 genes), fatty acid elongation (ko00062; 35 genes), and biosynthesis of unsaturated fatty acids (ko01040; 35 genes), were clearly represented. Specialized lipid pathways were also identified, such as ether lipid metabolism (ko00565; 32 genes), and three glycosphingolipid biosynthesis pathways—globo and iso-globo series (ko00603; 16 genes), ganglio series (ko00604; 7 genes), and lacto and neolacto series (ko00601; 5 genes) (Supplementary Tables S8–S9).

At the gene level, key enzymes involved in fatty acid biosynthesis were identified, including PT01_g01578 (β-ketoacyl-ACP synthase, K11262) and PT01_g01706 (malonyl-CoA-ACP transacylase, K01897). Fatty acid elongation and desaturation were supported by genes such as PT01_g03381 (3-ketoacyl-CoA synthase, K10248), PT01_g03561 (enoyl-CoA reductase, K10227), and PT01_g01956 (acyl-CoA oxidase, K00496). In the triacylglycerol (TAG) biosynthesis pathway, key enzymes such as DGAT1 (PT02_g06256, K11155) and PDAT1 (PT02_g07899, K13507) were identified as catalyzing the terminal acylation steps. Earlier and intermediate steps were facilitated by genes including GPAT (PT02_g06484, K22824), MGAT (PT01_g02271, K22825), and phosphatidate phosphatase (K01080), which also links phospholipid and TAG biosynthesis. Shared enzymes such as K00626 (PT01_g02514) and K00208 (PT01_g02497) were found to be involved in both fatty acid synthesis and additional lipid pathways, indicating functional overlap (Figure[3d; Supplementary Tables S8–S9; Supplementary Figure S5). Chromosomal mapping of lipid-related genes revealed a genome-wide distribution with evident clustering on chromosomes 2, 3, and 5 (Supplementary Figure S6). Functional categorization of these genes showed a predominance of unclassified enzymes, followed by oxidoreductases, acyltransferases, and hydrolases, consistent with the enzymatic diversity required for fatty acid biosynthesis, elongation, and remodeling.

### Orthologous gene clustering and cross-species conservation analysis

Orthologous gene clustering using high-confidence protein-coding genes from *P. tetragonolobus* and six related legumes—*Cajanus cajan*, *G. max*, *G. soja*, *Lotus japonicus*, *Vigna radiata*, and *Cicer arietinum*—identified a total of 21,342 orthogroups in *P. tetragonolobus*. *G. max* and *G. soja* showed the highest numbers of orthogroups with 35,359 and 35,026 clusters, respectively, while *C. arietinum* had the lowest (16,630). The orthogroup count for *P. tetragonolobus* was comparable to that of *C. cajan* (21,379) and *L. japonicus* (19,742) suggesting a similar degree of gene family complexity (Fig. 4a). An UpSet plot analysis revealed 14,365 orthogroups shared across all seven species, along with 12,769 shared exclusively between *G. max* and *G. soja*. Notably, *P. tetragonolobus* exhibited 1,358 species-specific orthogroups (Fig. 4b; Supplementary Fig. S5). A complementary jVenn diagram (Fig. 5) confirmed 928 orthogroups shared among all six species included in the analysis. Additionally, *P. tetragonolobus* possesses 1,358 species-specific orthogroups, while *L. japonicus* and *V. radiata* exhibit 1,285 and 719 unique orthogroups, respectively, underscoring both shared ancestry and lineage-specific innovations.

**Figure 4.**
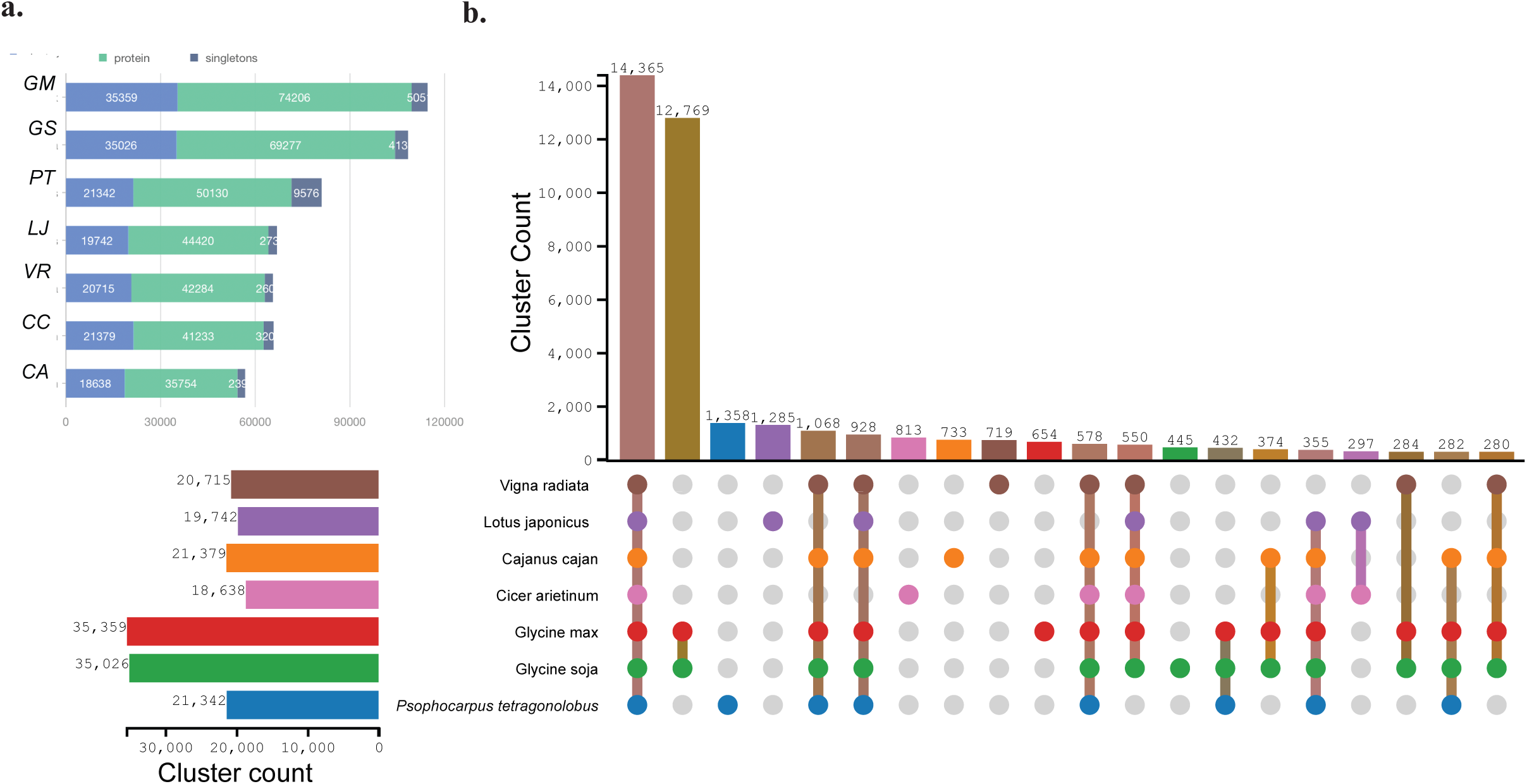
Comparative Analysis of Orthologous Gene Clusters in Legume Species. (a) Total number of orthologous gene clusters identified in seven legume species, including *Psophocarpus tetragonolobus (PT), Glycine max (GM), Glycine soja (GS), Cicer arietinum (CA), Cajanus cajan (CC), Lotus japonicus (LJ), and Vigna radiata (VR). G. max* and *G. soja* exhibit the highest number of gene clusters, while P. tetragonolobus shows a comparable number to C. cajan and V. radiata, indicating a substantial shared gene content among legumes. (b) Species-specific cluster counts highlight unique gene families in each species, with *P. tetragonolobus* possessing 1,358 unique clusters.

**Figure 5.**
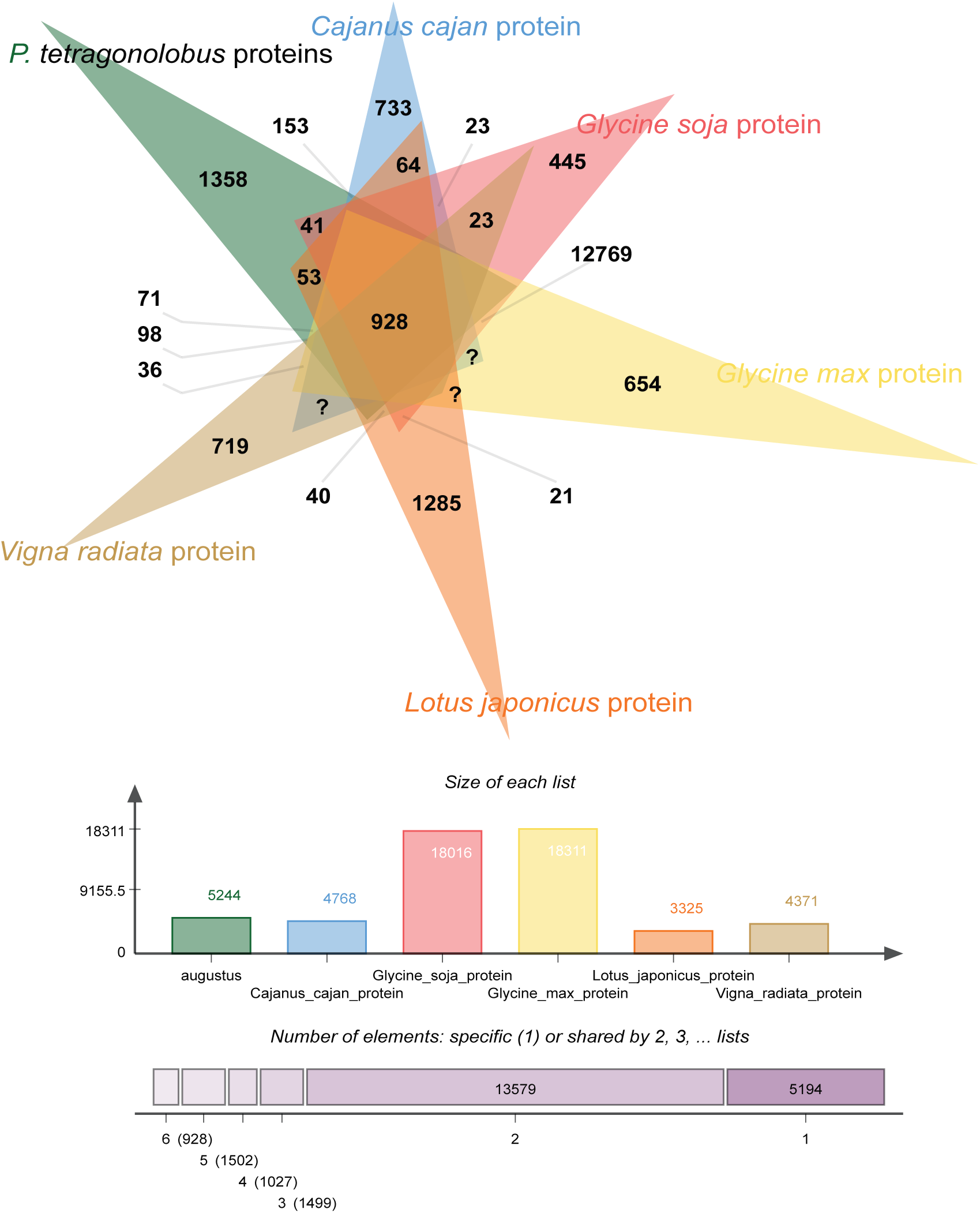
Orthologous Gene Family Intersections Among Six Legume Species. UpSet plot displaying the distribution of orthologous protein-coding gene families identified across six legumes: Psophocarpus tetragonolobus, Cajanus cajan, Glycine soja, Glycine max, Lotus japonicus, and Vigna radiata. The horizontal bars on the left represent the total number of orthologous gene families detected in each species. The vertical bars above the matrix indicate the number of gene families shared among the species combinations marked by filled circles in the matrix below. A total of 928 orthogroups are conserved across all six species (intersection with all circles filled), while P. tetragonolobus contains 1,358 species-specific orthogroups (intersection with only P. tetragonolobus filled), reflecting a substantial degree of lineage-specific gene family expansion. This pattern highlights both evolutionary conservation and divergence in gene content across legume genomes.

Gene family expansion and contraction analysis using CAFE indicated a contraction-dominated pattern in *P. tetragonolobus*, with 2,254 families lost and 91 gained—representing the highest contraction among the seven species examined (Fig. 6a). In contrast, the shared *G. max* –*G. soja* branch displayed 1,951 gene family gains and 167 losses. Individually, *G. max* gained 1,545 and lost 382 gene families, whereas *G. soja* gained 784 and lost 730. Intermediate turnover was noted in *C. cajan* and *V. radiata*, while *L. japonicus* and *C. arietinum* lost 1,301 and 1,415 gene families, respectively.

**Fig 6:**
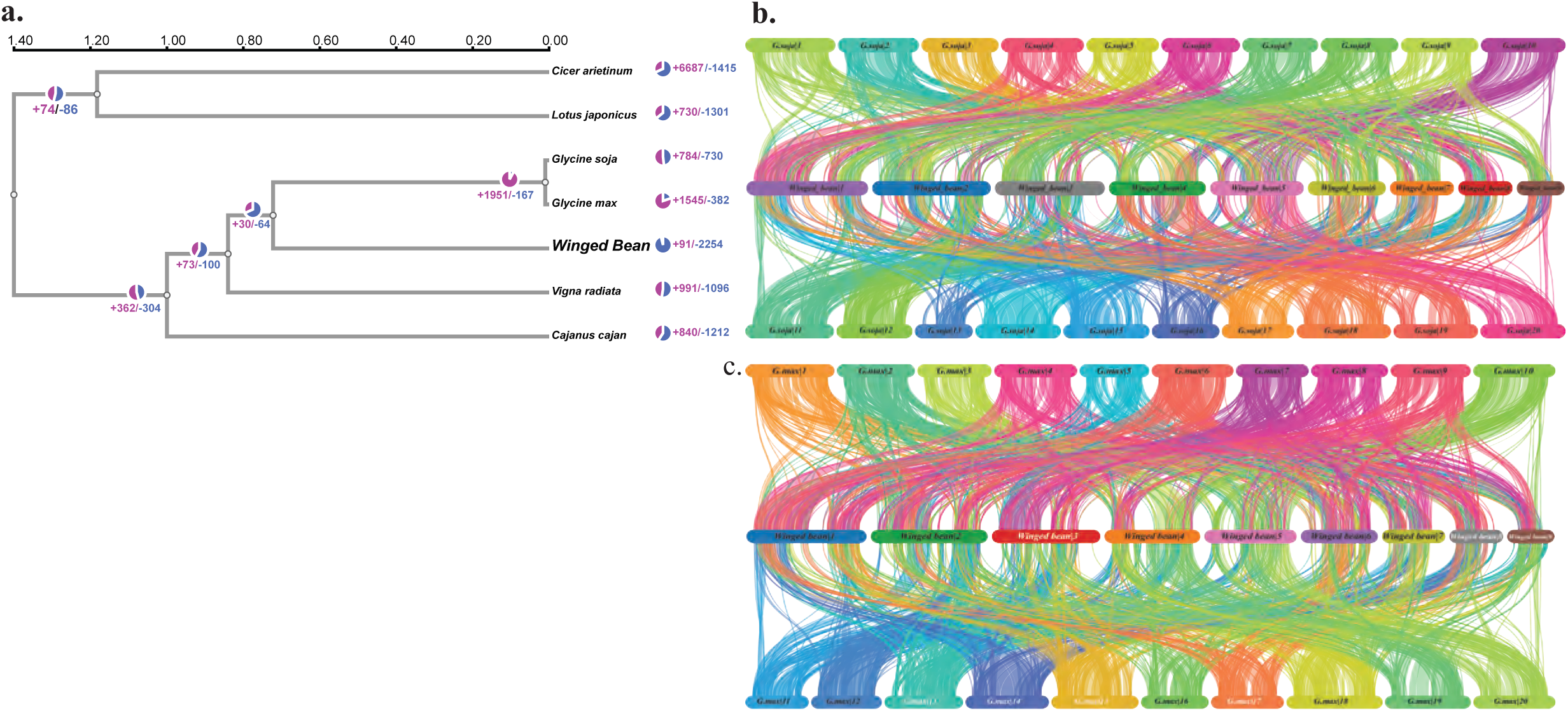
Expansion and Contraction of Gene Families and Synteny analysis. (a) Phylogenetic distribution of expanded (pink, +) and contracted (blue, –) gene families across eight legume species. The tree is based on conserved protein-coding genes. b–c) Chromosome-scale synteny plots comparing Psophocarpus tetragonolobus with (b) *Glycine soja* and (c) *Glycine max*. Extensive collinear blocks are observed, with a predominant 1:2 syntenic relationship, reflecting the additional whole-genome duplication events in Glycine species relative to the diploid *P. tetragonolobus*. This conserved macrosynteny highlights shared ancestry and structural divergence among legumes.

Genome-wide synteny analysis identified 40,162 syntenic blocks between *P. tetragonolobus* and three related legumes, comprising 1,504,765 homologous gene pairs. Comparative mapping with *G. max* revealed broad chromosome-scale collinearity, with most *P. tetragonolobus* chromosomes showing a 1:2 syntenic relationship. Syntenic blocks with *V. radiata* and *M. truncatula* were fewer and more fragmented, often localized to subtelomeric regions (Fig. 6b–c; Supplementary Fig. S7).

Orthogroup clustering of lipid-associated gene families across six legumes and *S. tuberosum* as an outlier revealed 26 clusters with species-specific distributions. *P. tetragonolobus* exhibited >5 gene copies in Cluster25923 and Cluster18019. Cluster14470, Cluster30006, and Cluster30478 showed higher gene copy numbers in *P. tetragonolobus* compared to other legumes, whereas Cluster526 and Cluster595 were present in only a few species (Fig. 7).

**Figure 7.**
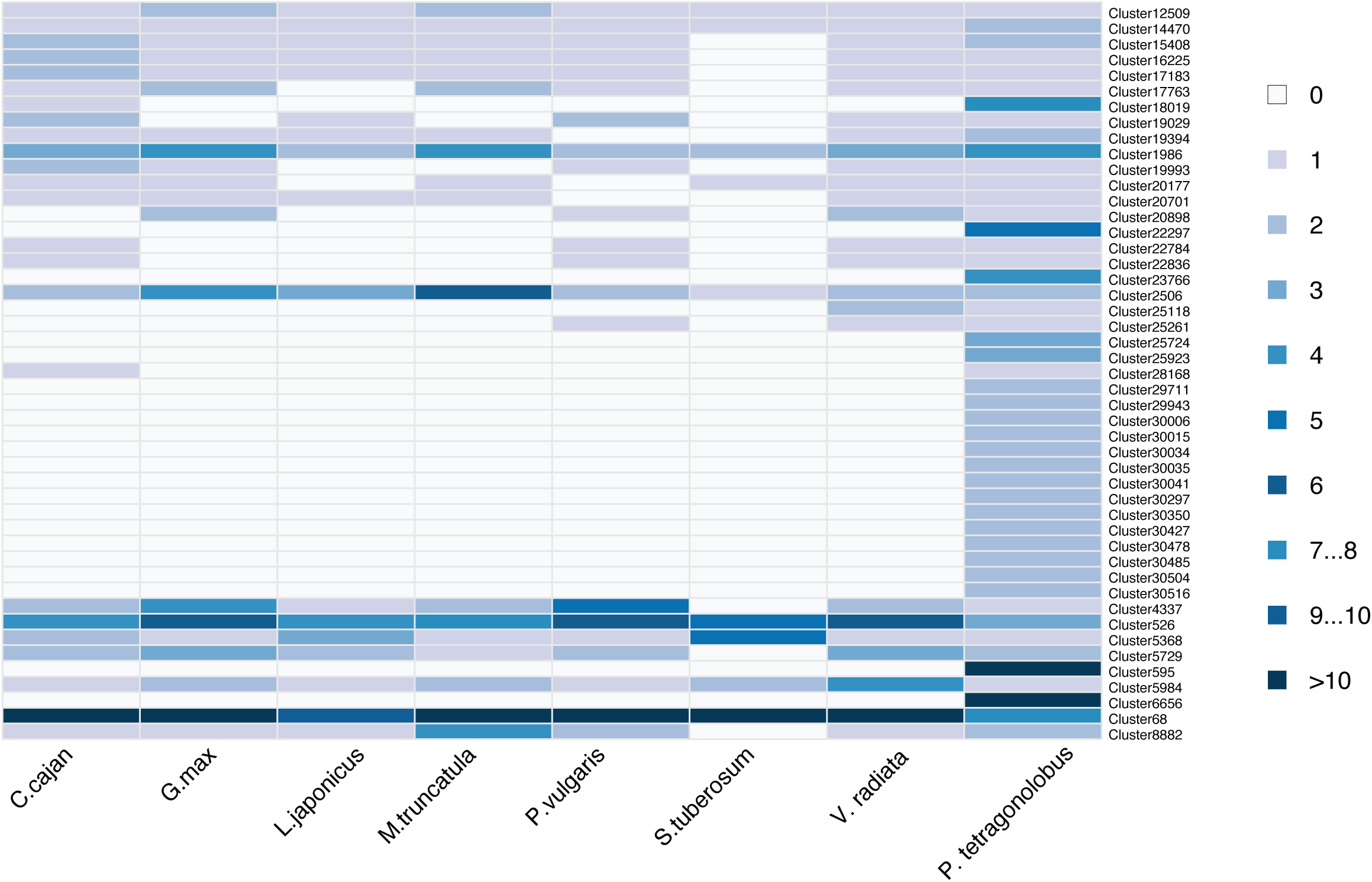
Heatmap showing species-wise distribution of gene family clusters associated with lipid metabolism. Each row represents a gene cluster identified through orthogroup clustering across nine plant species, while columns represent individual species, including P*sophocarpus tetragonolobus, Glycine max, Glycine soja, Phaseolus vulgaris, Vigna radiata, Lotus japonicus, Medicago truncatula, Solanum tuberosum, and Cajanus cajan. S.tuberosum* was included as an outgroup to explore potential metabolic convergence with *P. tetragonolobus*, which shares certain tuber-like storage traits.The color gradient indicates the number of genes from each species present in a given cluster, with darker shades reflecting higher gene counts. Notably, P. tetragonolobus exhibits enriched gene counts in several lipid-related clusters (e.g., Cluster14470, Cluster30006, Cluster30478), suggesting gene family expansion in key pathways involved in fatty acid biosynthesis, elongation, and storage lipid accumulation.

## Discussion

The winged bean genome presented here marks a significant leap in genomic characterization of underutilized legumes. With nearly complete chromosomal anchoring and high BUSCO completeness, the telomere-to-telomere (T2T) assembly offers one of the most complete legume references to date and provides foundational insights into genome structure, repeat composition, gene content, and evolutionary dynamics. Cytogenetic confirmation of a diploid, metacentric karyotype not only validates assembly accuracy but fills a longstanding gap in legume cytology for *P. tetragonolobus* in terms of chromosomal morphometric and ideotype characterization. Estimated flowcytometric genome size of ∼710.94 Mb, somewhat closer to an earlier report (782 Mb) by Bennet and Smith (1976)^11^, aligned well with the our assembled genome size of 697.69[Mb. However, our result contrasts earlier reports (1.22[Gbp/C^4^; 580.28[Mb^5^) where flowcytometric and assembled genome sizes from the same sample were not compared for accuracy. Compared to previous fragmented drafts (580.28 Mb in 46 scaffolds^5^), the present genome recovers ∼98.25% of the estimated size, successfully preserving complex repeat regions and structural motifs. This contiguity enables robust annotation and reliable mapping of functional genes, most notably, those involved in lipid metabolism, a trait central to winged bean’s agronomic appeal.

Interestingly, winged bean encodes over 750 lipid-related genes across 12 KEGG pathways, approaching counts observed in oil-rich polyploid legumes such as *G. max* (∼950 genes^12^) and *A. hypogaea* (∼900 genes^13,14^). Despite its diploid status and lack of recent whole-genome duplication (WGD), *P. tetragonolobus* maintains functional parity in key modules like triacylglycerol (TAG) biosynthesis, fatty acid elongation, and membrane lipid remodeling. Comprehensive annotation revealed 53,745 protein-coding genes in *P. tetragonolobus*, a number comparable to other major legumes such as *G. max* (∼46,430)^12^, *A. hypogaea* (∼66,469)^15^, *V. radiata* (∼42,289)^16^, and *P. sativum* (∼44,756)^17^, reaffirming the gene-rich and functionally diverse nature of this species. While *P. tetragonolobus* encodes fewer triacylglycerol (TAG) biosynthesis genes (n = 52) than *G. max* (65) or *A. hypogaea* (58)^18,19^, its seed oil content (∼16–21% of dry weight) approaches that of *G. max* (∼20%) and exceeds that of *P. vulgaris* (∼2%) and *C. arietinum* (∼6%)^20–22^, indicating that gene dosage alone does not dictate oil yield but rather, a tighter transcriptional regulation, enzyme efficiency, or pathway integration. However, further transcriptomic and metabolomic analyses are needed to confirm this hypothesis. TAG synthesis enzymes DGAT1 and PDAT1 are well-represented, while upstream reactions are supported by enzymes involved in acylation and phospholipid turnover. Chromosomal enrichment of lipid genes particularly on chromosomes 2, 3, and 5 suggests the presence of biosynthetic hotspots^23^ or legacy duplication zones retained through selective pressures^23,24^. The expansion of membrane lipid pathways, including ether lipids and glycosphingolipids, further highlights metabolic versatility, with implications for stress response, cellular stability, and niche adaptation. These traits, largely overlooked in legumes, merit further transcriptomic and metabolomic exploration.

Comparative genomics positions winged bean as a structurally stable genome undergoing contraction-dominated evolution. This contraction-biased pattern suggests lineage-specific genome streamlining or purifying selection, possibly driven by ecological specialization or the removal of redundant functions following ancient WGD^25,26^. This contraction-biased pattern suggests lineage-specific genome streamlining, potentially shaped by ecological specialization or reduced effective population size^27,28^. Although population-level genomic data were not generated in this study, recent work by Ho et al. (2024)^5^ analyzed 130 global accessions of winged bean and reported structured diversity, trait-associated QTLs, and domestication signatures affecting branching and pigmentation. These findings lend support to the hypothesis that gene family contraction in *P. tetragonolobus* may partially reflect selective constraints or bottlenecks in orphan crop evolution, particularly for agronomic ideotypes. The loss of over 2,200 gene families, coupled with preservation of 14,365 core orthogroups, reflects a streamlining trend, possibly arising from relaxed selection and domestication bottlenecks^29,30^. While speculative without population-level data, this trajectory mirrors patterns seen in *L. japonicus*^31^ and *C. arietinum*^32^, where gene loss accompanies ecological specialization.

Syntenic alignments with *G. max* reveal extensive chromosome-level conservation, with a 1:2 correspondence to *G. max* chromosomes consistent with paleopolyploidy in the *Glycine* lineage. The maintenance of macrosynteny despite gene family contraction suggests strong evolutionary constraints on core gene architecture in *P. tetragonolobus*. In contrast, comparisons with *V. radiata* and *M. truncatula* indicate greater divergence, underscoring winged bean’s unique evolutionary niche.

Taken together, our study indicates *P. tetragonolobus* as a powerful model for exploring trait co-regulation, compact genome evolution, and the genetic architecture underlying lipid accumulation. The insights provided here offer tangible routes for breeding climate-adaptive, protein- and oil-rich legumes and form the basis for gene discovery, pan-genomic surveys, and pathway engineering. In essence, *P. tetragonolobus* now emerges not only as a genetic resource but as a model for elevating orphan crops through high-resolution genomics, unlocking their value in sustainable food systems and nutrition security.

## Methods

### Plant material and Growth Conditions

The cultivar AKWB1 (IC0508386) was procured from the All India Coordinated Research Project on Potential Crops, ICAR-National Bureau of Plant Genetic Resources (NBPGR), New Delhi, India. To ensure genetic uniformity, AKWB1 was self-pollinated for three generations prior to the start of the experiment. Plants were grown under field conditions at the ICAR–Indian Institute of Agricultural Biotechnology (IIAB), Ranchi (23°16′27.6″N, 85°20′29.4″E). Leaf and root samples from 25-day-old plants were harvested for genome sequencing and cytogenetic analysis, and genome estimation respectively.

### Cytogenetic analysis and ideotyping

Actively growing root tips were pre-treated with 0.002M 8-hydroxyquinoline to arrest cells in metaphase, fixed in Carnoy’s solution (ethanol:acetic acid, 3:1), and stored in 70% ethanol. Chromosomes were stained using 4′,6-diamidino-2-phenylindole (DAPI) as described by Schweizer et al (2011)^33^ and visualized under a Zeiss Axiolab 5 fluorescence microscope. High-quality metaphase spreads (n = 7–10) were analyzed using DRAWID v0.26^34^ to measure arm lengths (long (q) and short (p)), total lengths, and arm ratios. Chromosomes were classified according to Baltisberger’s criteria, identifying morphology and ideotype structure^10^.

### Genome size estimation

Genome size was estimated by flow cytometry using *Oryza sativa* cv. Swarna (430 Mb) as an internal reference. Co-chopped leaf samples were stained with a propidium iodide (PI) buffer as described elsewhere^35^ and analyzed using BD FACS Canto equipped with a 488 nm laser and 582/42 nm band pass filter. Nuclei were acquired (n = 500–2000/sample) and dark incubated before exposure. G0/G1 peaks for both reference and sample showing a coefficient of variation (CV) below 5% were used for calculating mean fluorescence intensities which was then used to calculate genome size using the formula - *Sample genome size = Reference genome size x (Mean fluorescence intensity of sample / Mean fluorescence intensity of reference)*. Data were processed using FCS Express (DENOVO Software, USA).

### Genomic DNA isolation

Genomic DNA was extracted from young leaf tissue using the Blood & Cell Culture DNA Midi Kit (Qiagen, Cat No. 13343) following the manufacturer’s protocol. Quality and integrity were confirmed via agarose gel electrophoresis, NanoDrop® spectrophotometry, and FEMTO Pulse analysis. DNA concentrations were quantified in a Qubit 4.0 fluorometer using dsDNA HS Assay kit.

### SMRTbell library preparation and PacBio sequencing

Genomic DNA was sheared to a target size range of 12–18 kb using the Megaruptor 3 system with the Megaruptor-3 Shearing Kit. A total of 5 µg of gDNA in 60 µL (83.3 ng/µL) was loaded into hydropore tubes and sheared at a speed setting of 30. Sheared DNA was purified with 1× SMRTbell cleanup beads (PacBio) and eluted in 46 µL of elution buffer. Fragment size was confirmed on the FEMTO Pulse System. SMRTbell library construction was performed using the SMRTbell Express Template Prep Kit 3.0 following the manufacturer’s protocol. End-repair and A-tailing were carried out prior to ligation of non-barcoded SMRTbell adapters. Ligated products were cleaned with 1× SMRTbell cleanup beads and eluted in 40 µL of elution buffer. Nuclease treatment was performed to degrade unligated DNA and unbound adapters. Size selection was performed using AMPure PB beads to remove fragments <5[Kb. The final library was subjected to primer annealing and polymerase binding using the Sequel-II Binding Kit 3.2. Approximately 90 pM of the prepared library was loaded onto a SMRT Cell 8M and sequenced on the PacBio Sequel II platform (Nucleome Informatics Pvt. Ltd., Hyderabad, India) using circular consensus sequencing (CCS/HiFi) mode.

### Illumina library preparation and sequencing

The high-quality genomic DNA extracted from AKWB1 was also used in Illumina whole-genome sequencing to ensure consistency across platforms. A total of 500 ng of DNA was used to construct the library using the KAPA HyperPlus Kit, following the manufacturer’s protocol. Library concentration was quantified using a Qubit 4.0 Fluorometer with the dsDNA HS Assay Kit. Fragment size distribution was evaluated on the Agilent-2100 Bioanalyzer using the High Sensitivity DNA Kit. Sequencing was performed on the Illumina NovaSeq 6000 platform (Nucleome Informatics Pvt. Ltd., Hyderabad, India) using an S4 flow cell and 2[×[150[bp paired-end chemistry.

### HiC Sequencing

Hi-C libraries were prepared using the Proximo Hi-C Plant Kit following the manufacturer’s protocol. Approximately 1 g of leaf tissue was used. Briefly, tissue was crosslinked, lysed, and chromatin was fragmented, followed by proximity ligation to capture spatially adjacent DNA fragments. Crosslinks were reversed and DNA was purified for library construction via adapter ligation and PCR amplification. Final Hi-C libraries were quantified using a Qubit 4.0 fluorometer with the DNA High Sensitivity assay kit. Fragment size was analyzed using an Agilent-2100 Bioanalyzer. Hi-C libraries with fragment sizes in the range of 400–500 bp were sequenced on an Illumina NovaSeq 6000 platform (S4 flow cell, 2[×[150 bp paired-end reads) at Nucleome Informatics Pvt. Ltd., Hyderabad, India.

### Optical mapping

Ultra-high molecular weight (UHMW) genomic DNA was extracted from plant tissue using the Bionano Prep™ Plant Tissue DNA Kit following the manufacturer’s protocol. Ultra-high molecular weight (UHMW) DNA (750 ng) was labeled using the Direct Label and Stain (DLS) protocol with the Bionano Prep DLS DNA Labeling Kit, following the manufacturer’s instructions. Labeled DNA was purified, loaded onto a Saphyr chip, and imaged using the Saphyr optical mapping system (Nucleome Informatics Pvt. Ltd., Hyderabad, India).

### Chromosome scale genome assembly

High-quality subread data were generated using the PacBio Sequel II platform, and high-fidelity (HiFi) reads were obtained by processing the subreads with the Circular Consensus Sequencing (CCS) algorithm^36^. De novo genome assembly was performed using Hifiasm v0.25.0-r726^37,38^. Base-level accuracy and overall quality of the draft genome assembly were enhanced by aligning Illumina paired-end short reads using BWA-MEM v0.7.18^39^. Small indels and SNPs of the draft genome were corrected using Pilon v1.24^40^. Chromosome-scale scaffolding was performed by integrating BioNano optical maps using the Hybrid Scaffold pipeline, followed by Hi-C data processing with Juicer v180419^41^ and manual curation using Juicebox Assembly Tools^42^. The completeness of the draft genome assembly was assessed using BUSCO v5.8.2^43^ .

### Comprehensive identification of repetitive elements and gene prediction

Repetitive elements were identified, annotated, and classified using RepeatModeler v2.9.5^44^. Transposable elements (TEs) were further characterized using the De Novo TE Annotator (EDTA) pipeline^45^ with default parameters. EDTA integrates homology-based and structural approaches to detect and classify diverse TEs, including LTR retrotransposons, DNA transposons, and other interspersed repeats. The BRAKER3 pipeline^46^, an integrated de novo and homology-based gene annotation approach, was used to predict genome-wide protein-coding genes. The pipeline trained GeneMark-ETP and AUGUSTUS, using RNA-seq evidence and homology support from both *G. max* (as the closest annotated legume reference) and the viridiplantae_odb10 ortholog set. This dual strategy enabled precise gene model prediction tailored to legume genomes^47^.

### Lipid metabolic pathway analysis

Lipid metabolism-related genes were annotated by mapping KEGG Orthology (KO) and Enzyme Commission (EC) identifiers using EggNOG-mapper v5^48^. Genes were assigned KEGG pathways, including those involved in fatty acid biosynthesis (ko00061), elongation (ko00062), and triacylglycerol (TAG) assembly (ko00561), enabling identification of key biosynthetic modules^49^. Pathway visualizations were generated using the R packages *Pathview* ^50^ and *PathfinderR* ^51^, with mapped genes overlaid onto KEGG reference maps. Functional annotation was refined using InterProScan to identify conserved domains and motifs, with Gene Ontology (GO) terms assigned based on InterPro and EggNOG annotations, supported by NCBI NR, Pfam, Swiss-Prot, TrEMBL, and eggNOG database searches^52^. GO enrichment analysis was conducted using *clusterProfiler* v4.4.4^53^, and results were visualized as ontology-classified dot plots using *ggplot2* v3.4.2^54^. The chromosomal distribution of lipid metabolism-related genes was visualized using the R package karyoploteR v1.28.0^55^. Lipid metabolism genes were curated from genome annotations using KEGG Orthology and EC identifiers, and classified into functional categories such as acyltransferases, desaturases, and lipases. TAG biosynthesis genes were filtered from the annotated genome using a combination of KEGG pathway mapping (ko00561) and domain-based annotations from InterProScan and EggNOG-mapper. Only genes with confirmed enzymatic roles in the Kennedy pathway and TAG remodeling were included in the final count.

### Comparative genomics

Comparative genomic analysis was conducted to identify orthologous gene clusters among *P. tetragonolobus* and six legume species: *G. max*, *G. soja*, *V. radiata*, *C. cajan*, *L. japonicus*, and *C. arietinum* using the OrthoVenn3 platform (https://orthovenn3.bioinfotoolkits.net/home)^56^. Predicted protein sequences were subjected to all-against-all BLASTP^57^ with an E-value cutoff of 1e-5, and orthogroups were defined using the Markov Cluster Algorithm (MCL) with an inflation value of 1.5. Functional conservation and lineage-specific divergence were inferred from OrthoVenn3 outputs^56^ Syntenic relationships between *P. tetragonolobus* and other four legume species—*G. max*, *G. soja*, *V. radiata*, and *M. truncatula* were investigated using MCScanX^58^. Protein-coding sequences were aligned using BLASTP. Gene coordinates were formatted in MCScanX-compatible GFF format, and collinear blocks were detected using default parameters (≥5 collinear genes per block; ≤25 intervening genes). The resulting *.collinearity* files were converted to JCVI-compatible format using the *jcvi.compara.synteny* module, and multi-genome synteny maps were generated using the *graphics.karyotype* and *graphics.synteny* modules of JCVI^59^, allowing chromosome-scale visualization of conserved genomic blocks and structural rearrangements across species.

### Data restructuring and visualization

All raw outputs from annotation, orthology, enrichment, and synteny analyses were programmatically parsed and reformatted into structured data frames using base R and Python scripts. These scripts were adapted to meet the input requirements of downstream tools and plotting libraries.

## Data Availability

The reference genome assembly of *Psophocarpus tetragonolobus* has been submitted to the NCBI under BioProject PRJNA1169650 with accession number JBMOEB000000000

## Code availability

No custom algorithm development was required, and all transformations followed standard scripting logic. Code is available upon request.

## Acknowledgements

The authors express their immense gratitude to Nucleome Informatics Pvt. Limited, Hyderabad for their invaluable support in this research. We also acknowledge Dr. Prakit Somta, Associate Professor, Department of Agronomy, Faculty of Agriculture at Kamphaeng Saen, Kasetsart University, Nakhon Pathom 73140, Thailand, for kindly sharing the winged bean linkage map. Although the shared resource was initially considered for use in our study, we ultimately succeeded in generating a high-quality genome assembly independently.

## Author Information

**ICAR-Indian Institute of Agricultural Biotechnology, Ranchi - 834003, Jharkhand (India)**

Kishor U. Tribhuvan, Nikhil Kumar Singh, Binay Kumar Singh, Avinash Pandey, Sudhir Kumar, Sujit Kumar Bishi, Kanaka K. K, Tanmaya Kumar Sahu, A. Pattanayak, Vijai Pal Bhadana, Sujay Rakshit

## Contributions

K.U.T. - Planning of experiment, genome size estimation, sequencing, data analysis and writing of manuscript, N.K.S. - Data analysis, writing of manuscript, B.K.S. - Planning of experiment, genome size estimation, sequencing, writing manuscript, A.P. - Data Analysis and writing of manuscript, S.K. - Generation of plant material, writing of manuscript, S.K.B. - Manuscript writing, K.K.K.- Data interpretation and writing of manuscript, T.K.S. - Data Analysis and writing of manuscript, A.Pt. - Coordination of project, Manuscript Editing, V.P.B.- Coordination of project, Manuscript Editing, S.R.- Coordination of project, Interpretation, writing of manuscript

## Supplementary information

Supplementary Figures. S1–S7, along with their corresponding supplementary figure legends, and Supplementary Tables S1–S10.

