## Supplementary table S1 for "Chromosome-Scale, Telomere-to-Telomere Assembly of Winged Bean (*Psophocarpus tetragonolobus* (L.) DC.) Genome Reveals Lipid Biosynthetic Hotspots and Gene Family Dynamics"

**Supplementary Table S1:** Chromosome length, long arm and short arm length, and arm ratio of winged bean chromosomes.

| **Chromosome** | **Homolog** | **Total Length (µm)** | **Short Arm (q) (µm)** | **Long Arm (p) (µm)** | **Arm Ratio (p/q)** | **SEM (Total)** | **Chromosome Type** |
| --- | --- | --- | --- | --- | --- | --- | --- |
| 1 | A | 182.08 | 65.20 | 116.88 | 1.79 |  | Metacentric |
|  | B | 200.74 | 82.10 | 118.64 | 1.45 |  |  |
|  | **Average** | **191.41** | **73.65** | **117.76** | **1.60** | **9.33** |  |
| 2 | A | 169.01 | 69.51 | 99.50 | 1.43 |  | Metacentric |
|  | B | 189.50 | 74.96 | 114.54 | 1.53 |  |  |
|  | **Average** | **179.26** | **72.24** | **107.02** | **1.48** | **10.25** |  |
| 3 | A | 152.12 | 69.13 | 82.99 | 1.20 |  | Metacentric |
|  | B | 167.22 | 76.26 | 90.96 | 1.19 |  |  |
|  | **Average** | **159.67** | **72.70** | **86.98** | **1.20** | **7.55** |  |
| 4 | A | 145.19 | 60.99 | 84.20 | 1.38 |  | Metacentric |
|  | B | 166.39 | 78.78 | 87.61 | 1.11 |  |  |
|  | **Average** | **155.79** | **69.89** | **85.91** | **1.23** | **10.60** |  |
| 5 | A | 147.50 | 64.53 | 82.97 | 1.29 |  | Metacentric |
|  | B | 154.65 | 74.16 | 80.49 | 1.09 |  |  |
|  | **Average** | **151.08** | **69.35** | **81.73** | **1.18** | **3.58** |  |
| 6 | A | 124.57 | 55.21 | 69.36 | 1.26 |  | Metacentric |
|  | B | 129.81 | 57.43 | 72.38 | 1.26 |  |  |
|  | **Average** | **127.19** | **56.32** | **70.87** | **1.26** | **2.62** |  |
| 7 | A | 118.95 | 50.29 | 68.66 | 1.37 |  | Metacentric |
|  | B | 119.96 | 50.94 | 69.02 | 1.35 |  |  |
|  | **Average** | **119.46** | **50.62** | **68.84** | **1.36** | **0.51** |  |
| 8 | A | 115.19 | 50.02 | 65.17 | 1.30 |  | Metacentric |
|  | B | 123.98 | 61.00 | 62.98 | 1.03 |  |  |
|  | **Average** | **119.59** | **55.51** | **64.08** | **1.15** | **4.40** |  |
| 9 | A | 109.33 | 52.95 | 56.38 | 1.06 |  | Metacentric |
|  | B | 116.11 | 54.16 | 61.95 | 1.14 |  |  |
|  | **Average** | **112.72** | **53.56** | **59.17** | **1.10** | **3.39** |  |

Winged bean is a diploid crop species, with each somatic cell containing a pair of homologous chromosomes.
