## Supplementary table S2 for "Chromosome-Scale, Telomere-to-Telomere Assembly of Winged Bean (*Psophocarpus tetragonolobus* (L.) DC.) Genome Reveals Lipid Biosynthetic Hotspots and Gene Family Dynamics"

**Supplementary Table S2:** Summary of DNA sequencing data

| **Read type** | **Number of clean reads / subreads** | **Insert size (bp)** | **Total data (Gb)** | **Read length /**  **Mean subread length (N50) (bp)** | **Sequence coverage***  **(X)** |
| --- | --- | --- | --- | --- | --- |
| **Illumina pair-end reads** | 2061949860 | 400 | 311.35 | 150 | 338X |
| **Hi-C pair end reads** | 301912198 | 700 | 45.286 | - | 65X |
| **Pacbio subreads** | 11993641 | 20 KB | 990.683 | 182250 | 1415X |
| **Optical Mapping** | No. Of molecules 11237484 |  | Total DNA (>= 20kbp) 652.29 Gbp  N50 (>= 20 kbp) 78.75 kbp  Total DNA (>= 150kbp)76.21 Gbp  N50 (>= 150kbp) 184.13 kbp  Total DNA (>= 150kbp and min sites >=9) 68.48 Gbp  N50 (>= 150kbp and min sites >=9) 184.88 kbp | - | 34.76X |

*Depth was calculated under the estimate of a genome size of 710 Mb.
