## Supplementary table S3 for "Chromosome-Scale, Telomere-to-Telomere Assembly of Winged Bean (*Psophocarpus tetragonolobus* (L.) DC.) Genome Reveals Lipid Biosynthetic Hotspots and Gene Family Dynamics"

**Supplementary Table S3**: Progressive Improvement in Winged Bean Genome Assembly Using PacBio HiFi, BioNano, and Hi-C Technologies

| **Assembly statistics** | **PacBio HiFi** | **PacBio HiFi Haplotypic duplication purged** | **PacBio + BioNano** | **PacBio + BioNano + Hi-C** |
| --- | --- | --- | --- | --- |
| **Total genome size** | 862231031 | 716851204 | 697677570 | 697,685,870 |
| **Total number of fragments** | 1294 | 165 | 59 | 15 |
| **Largest fragment size** | 46,951,877 | 46,951,877 | 88,789,861 | 111,651,818 |
| **L50 (Number)** | 18 | 13 | 9 | 4 |
| **N50 (Size)** | 14,621,422 | 16,936,191 | 19,362,744 | 85,977,384 |
| **L90 (Number)** | 87 | 43 | 32 | 8 |
| **N90 (Size)** | 1,107,953 | 4,302,283 | 5,996,666 | 41,789,641 |
| **Total number of Gaps** | 0 | 0 | 65 | 111 |
| **Ambiguous Bases (N) per 100kbp** | 0 | 0 | 26.83 | 27.47 |
| **BUSCO^*^ (%)** | C:97.2% [S:88.7%, D:8.5%], F:0.3%, M:2.5%, n:5366 | C:97.2% [S:93.2%, D:4.0%], F:0.3%, M:2.5%, n:5366 | C:97.0% [S:93.3%, D:3.7%], F:0.3%, M:2.7%, n:5366 | C:97.61% [S:94.5%, D:3.1%], F:0.4%, M:1.9%, n:5366 |
