## Supplementary table S4 for "Chromosome-Scale, Telomere-to-Telomere Assembly of Winged Bean (*Psophocarpus tetragonolobus* (L.) DC.) Genome Reveals Lipid Biosynthetic Hotspots and Gene Family Dynamics"

**Supplementary Table S4:** Chromosome-wise distribution of the final *Psophocarpus tetragonolobus* genome assembly

| **Chromosome** | **Length (bp)** |
| --- | --- |
| CHR1 | 111,651,818 |
| CHR2 | 108,313,712 |
| CHR3 | 99,375,205 |
| CHR4 | 85,977,384 |
| CHR5 | 82,569,713 |
| CHR6 | 66,280,345 |
| CHR7 | 51,943,675 |
| CHR8 | 41,789,641 |
| CHR9 | 34,859,452 |
| **Subtotal (anchored)** | **682,760,945** |
| Unplaced scaffold 1 | 8,563,885 |
| Unplaced scaffold 2 | 2,712,448 |
| Unplaced scaffold 3 | 1,387,345 |
| Unplaced scaffold 4 | 1,303,631 |
| Unplaced scaffold 5 | 825,561 |
| Unplaced scaffold 6 | 132,055 |
| **Subtotal (unplaced)** | **14,924,925** |
| **Total assembly size** | **697,685,870** |

Chromosome-wise distribution of the *P. tetragonolobus* genome after Hi-C-based scaffolding. A total of 682.76 Mb (97.73%) of the assembled genome was anchored to nine pseudochromosomes, corresponding to the haploid chromosome number of the species. The remaining 14.92 Mb (2.14%) was retained in six unplaced scaffolds.
