## Supplementary table S5 for "Chromosome-Scale, Telomere-to-Telomere Assembly of Winged Bean (*Psophocarpus tetragonolobus* (L.) DC.) Genome Reveals Lipid Biosynthetic Hotspots and Gene Family Dynamics"

**Supplementary Table S5**: Distribution and Abundance of Transposable Elements and Repeats in the *P. tetragonolobus* Genome

| Sr. No | Repeat Class | Number of Elements | Length (bp) | Percentage of Genome (%) |
| --- | --- | --- | --- | --- |
| 1. | **Total interspersed repeats** | **202786** | **409656529** | **60.00** |
|  | LINEs | 1448 | 220701348 | 32.33 |
|  | LTR elements | 260 | 77151768 | 11.29 |
|  | DNA transposons | 389 | 63584421 | 9.31 |
|  | Unclassified | 316 | 48218992 | 7.06 |
| 2. | **Small RNA-associated repeats** | **5782** | **386402979** | **56.59** |
| 3. | **Satellites** | **49** | **3486** | **0** |
| 4. | **Simple sequence repeats (SSRs)** | **306253** | **28631307** | **4.19** |
| 5. | **Low complexity regions** | **88142** | **16238136** | **2.38** |
