## Supplementary table S7 for "Chromosome-Scale, Telomere-to-Telomere Assembly of Winged Bean (*Psophocarpus tetragonolobus* (L.) DC.) Genome Reveals Lipid Biosynthetic Hotspots and Gene Family Dynamics"

**Supplementary Table S7:** Functional annotation of the *P. tetragonolobus* genes

| **Annotation Type** | **Annotated Genes** | **Percentage** |
| --- | --- | --- |
| NR (BLAST) | 39909 | 74.3 |
| InterPro (any match) | 53712 | 99.94 |
| Pfam | 25223 | 46.9 |
| GO Slim | 28451 | 52.9 |
| EggNOG | 34607 | 70.01 |
| GO (EggNOG) | 15462 | 31.28 |
| EC | 6451 | 13.05 |
| KEGG Orthologs (KO) | 14089 | 28.50 |
| KEGG Pathway | 8852 | 17.91 |
| KEGG Module | 14089 | 28.50 |
| KEGG Reaction | 1569 | 3.17 |
| KEGG Reaction Class | 731 | 1.48 |
| KEGG Transporters (TC) | 28072 | 56.79 |
