## Supplementary material for "Chromosome-Scale, Telomere-to-Telomere Assembly of Winged Bean (*Psophocarpus tetragonolobus* (L.) DC.) Genome Reveals Lipid Biosynthetic Hotspots and Gene Family Dynamics": Table 1

**Table 1:** Global statistics of *Psophocarpus tetragonolobus* genome assembly and annotation

| **Parameter** | **Value** |
| --- | --- |
| Genome size estimation (Mb) | 710.94 |
| Assembled genome size (bp) | 697,685,870 |
| Total number of scaffolds | 15 |
| Largest scaffold (bp) | 111,651,818 |
| Scaffold N50 (bp) | 85,977,384 |
| Scaffold L50 | 4 |
| Scaffold N90 (bp) | 41,789,641 |
| Scaffold L90 | 8 |
| Total number of gaps | 111 |
| Ambiguous bases (N) per 100 kbp | 27.47 |
| GC content (%) | 31.08 |
| BUSCO completeness (%) | 95.7 |
| Total length of repetitive elements (bp) | 409,656,529 |
| Repeat content (%) | 60.00 |
| LTR retrotransposon density (%) | 11.30 |
| Microsatellite repeat density (%) | 4.19 |
| Total number of annotated genes | 53,746 |
