## Supplementary figure S1 for "Chromosome-Scale, Telomere-to-Telomere Assembly of Winged Bean (*Psophocarpus tetragonolobus* (L.) DC.) Genome Reveals Lipid Biosynthetic Hotspots and Gene Family Dynamics"

### Slide 1
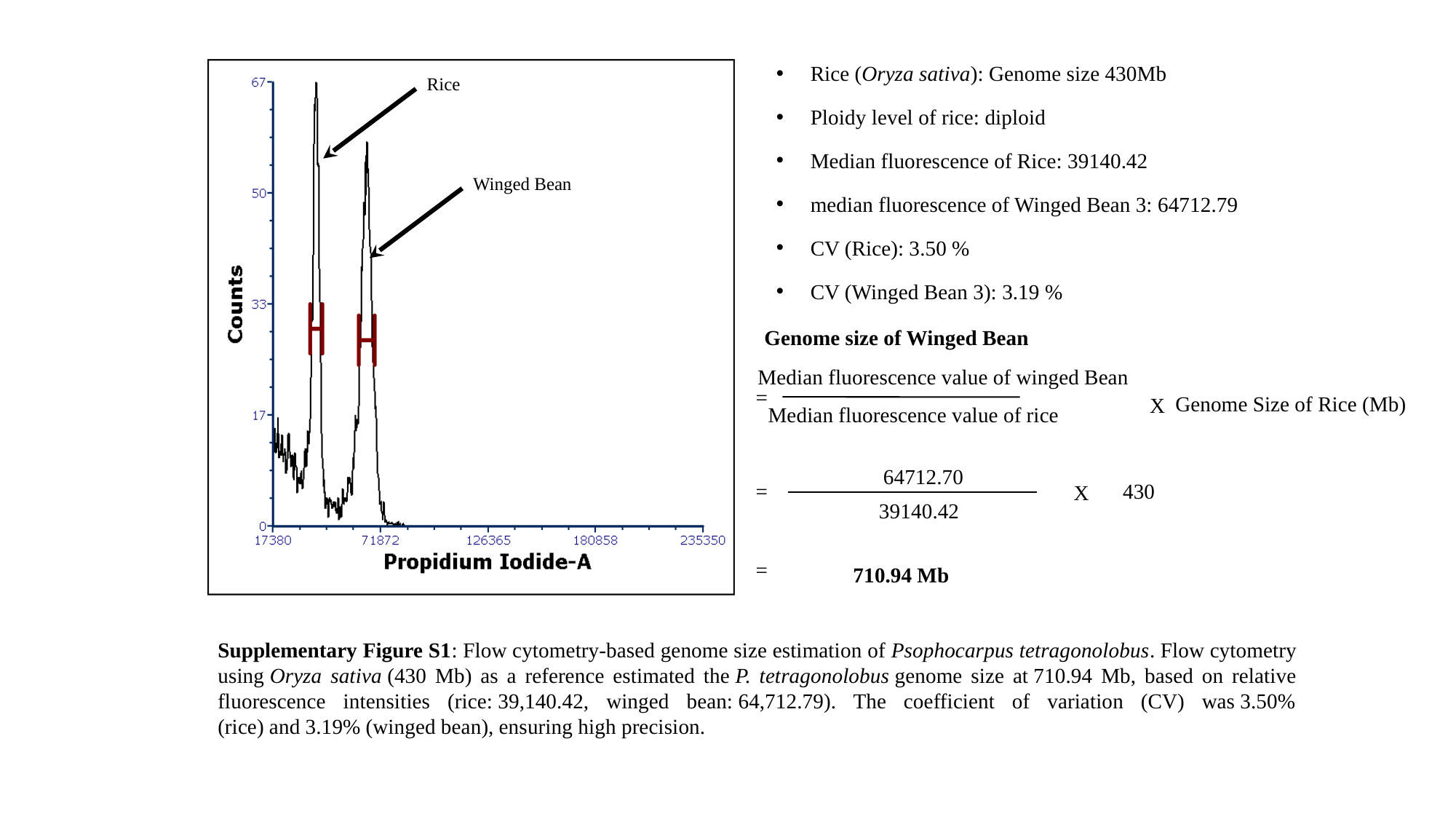

Rice (Oryza sativa): Genome size 430Mb
Ploidy level of rice: diploid
Median fluorescence of Rice: 39140.42
median fluorescence of Winged Bean 3: 64712.79
CV (Rice): 3.50 %
CV (Winged Bean 3): 3.19 %
Rice
Winged Bean
Genome size of Winged Bean
Median fluorescence value of winged Bean
=
Genome Size of Rice (Mb)
X
Median fluorescence value of rice
64712.70
=
430
X
39140.42
=
710.94 Mb
Supplementary Figure S1: Flow cytometry-based genome size estimation of Psophocarpus tetragonolobus. Flow cytometry using Oryza sativa (430 Mb) as a reference estimated the P. tetragonolobus genome size at 710.94 Mb, based on relative fluorescence intensities (rice: 39,140.42, winged bean: 64,712.79). The coefficient of variation (CV) was 3.50% (rice) and 3.19% (winged bean), ensuring high precision.
