## Supplementary figure S2 for "Chromosome-Scale, Telomere-to-Telomere Assembly of Winged Bean (*Psophocarpus tetragonolobus* (L.) DC.) Genome Reveals Lipid Biosynthetic Hotspots and Gene Family Dynamics"

### Slide 1
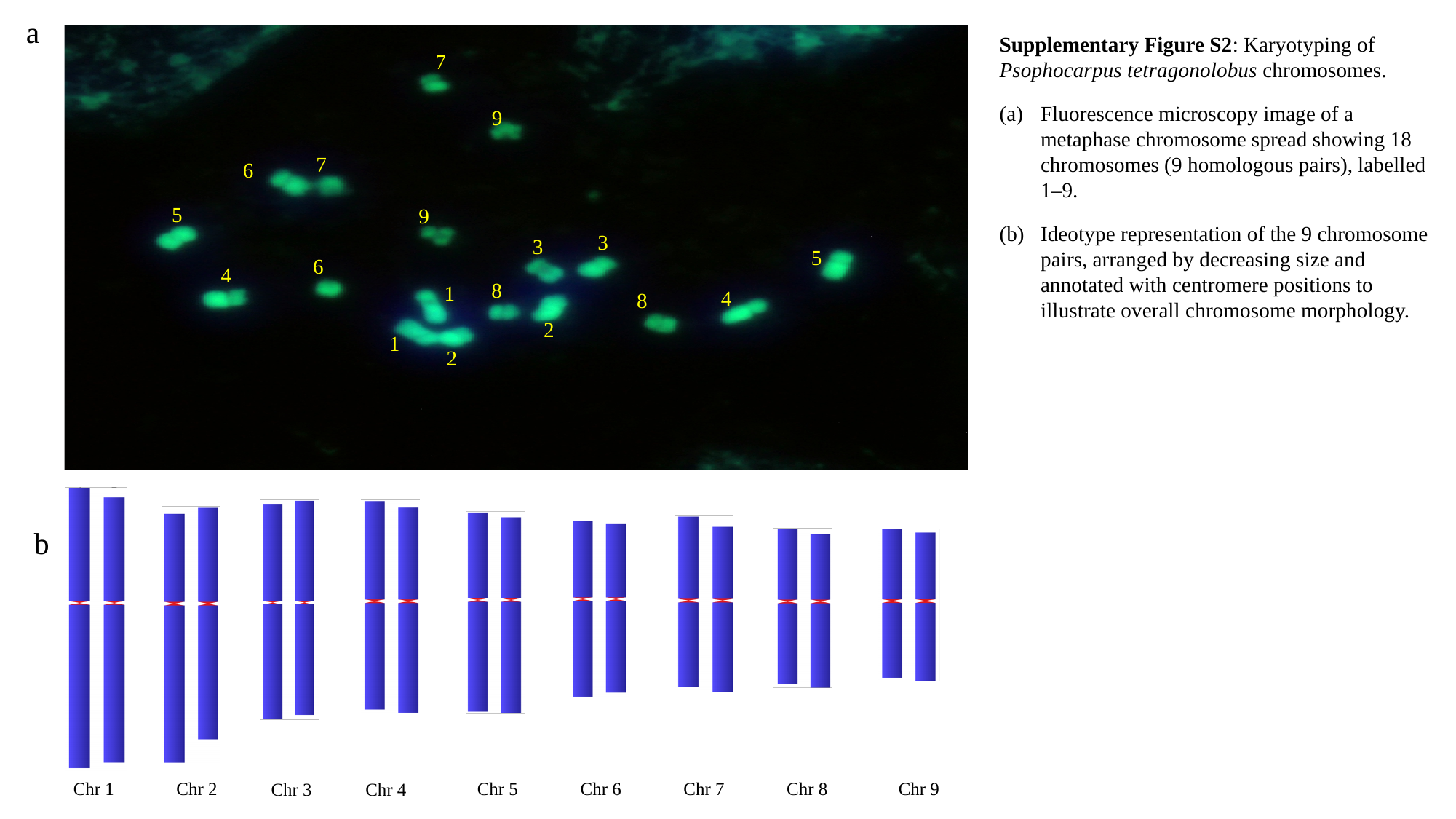

a
Supplementary Figure S2: Karyotyping of Psophocarpus tetragonolobus chromosomes.
Fluorescence microscopy image of a metaphase chromosome spread showing 18 chromosomes (9 homologous pairs), labelled 1–9.
Ideotype representation of the 9 chromosome pairs, arranged by decreasing size and annotated with centromere positions to illustrate overall chromosome morphology.
7
9
7
6
5
9
3
3
5
6
4
8
1
4
8
2
1
2
Chr 1
Chr 2
Chr 5
Chr 6
Chr 7
Chr 8
Chr 9
Chr 3
Chr 4
b
