## Supplementary figure S3 for "Chromosome-Scale, Telomere-to-Telomere Assembly of Winged Bean (*Psophocarpus tetragonolobus* (L.) DC.) Genome Reveals Lipid Biosynthetic Hotspots and Gene Family Dynamics"

### Slide 1
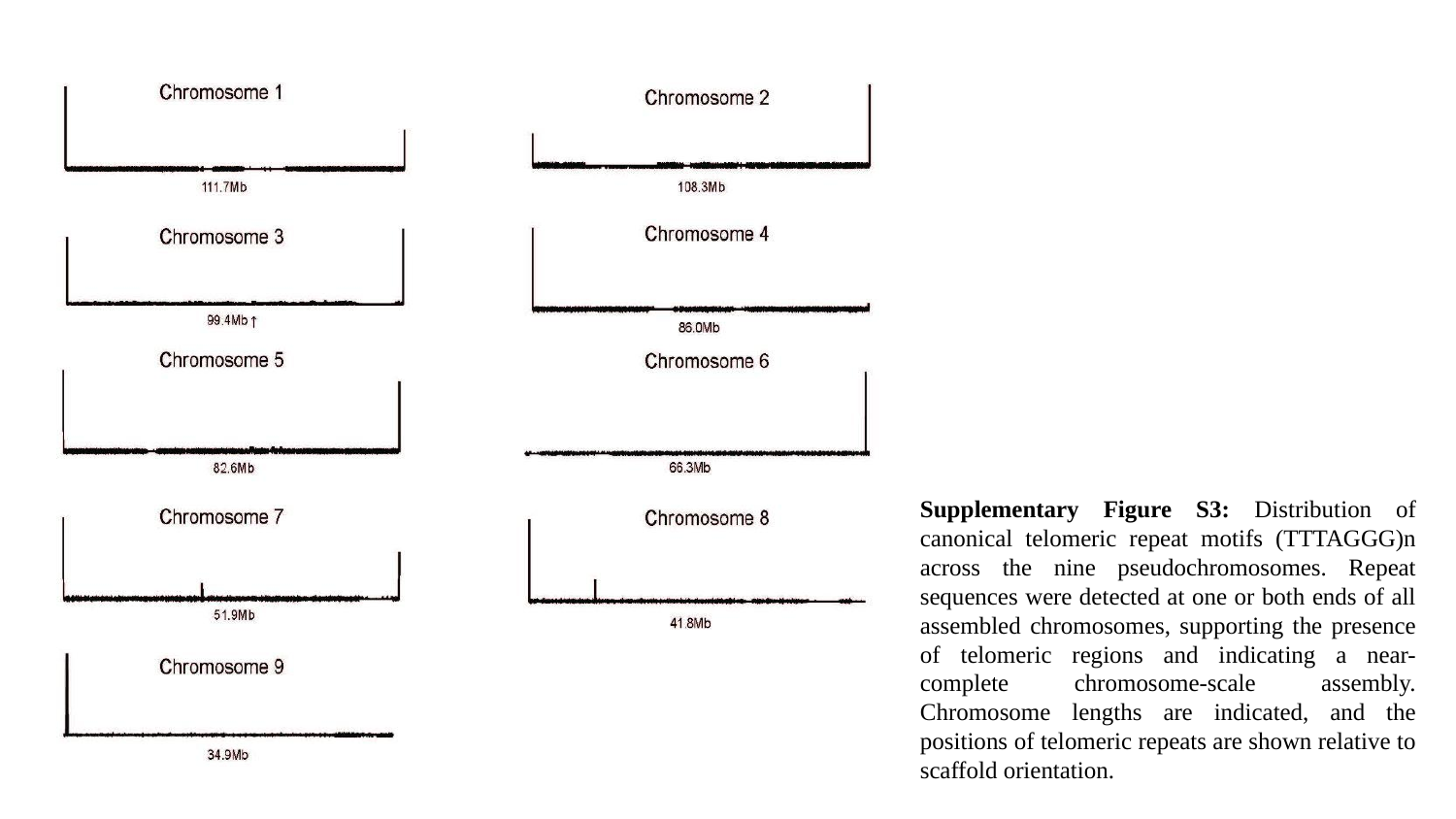

Supplementary Figure S3: Distribution of canonical telomeric repeat motifs (TTTAGGG)n across the nine pseudochromosomes. Repeat sequences were detected at one or both ends of all assembled chromosomes, supporting the presence of telomeric regions and indicating a near-complete chromosome-scale assembly. Chromosome lengths are indicated, and the positions of telomeric repeats are shown relative to scaffold orientation.
