## Supplementary figure S4 for "Chromosome-Scale, Telomere-to-Telomere Assembly of Winged Bean (*Psophocarpus tetragonolobus* (L.) DC.) Genome Reveals Lipid Biosynthetic Hotspots and Gene Family Dynamics"

### Slide 1
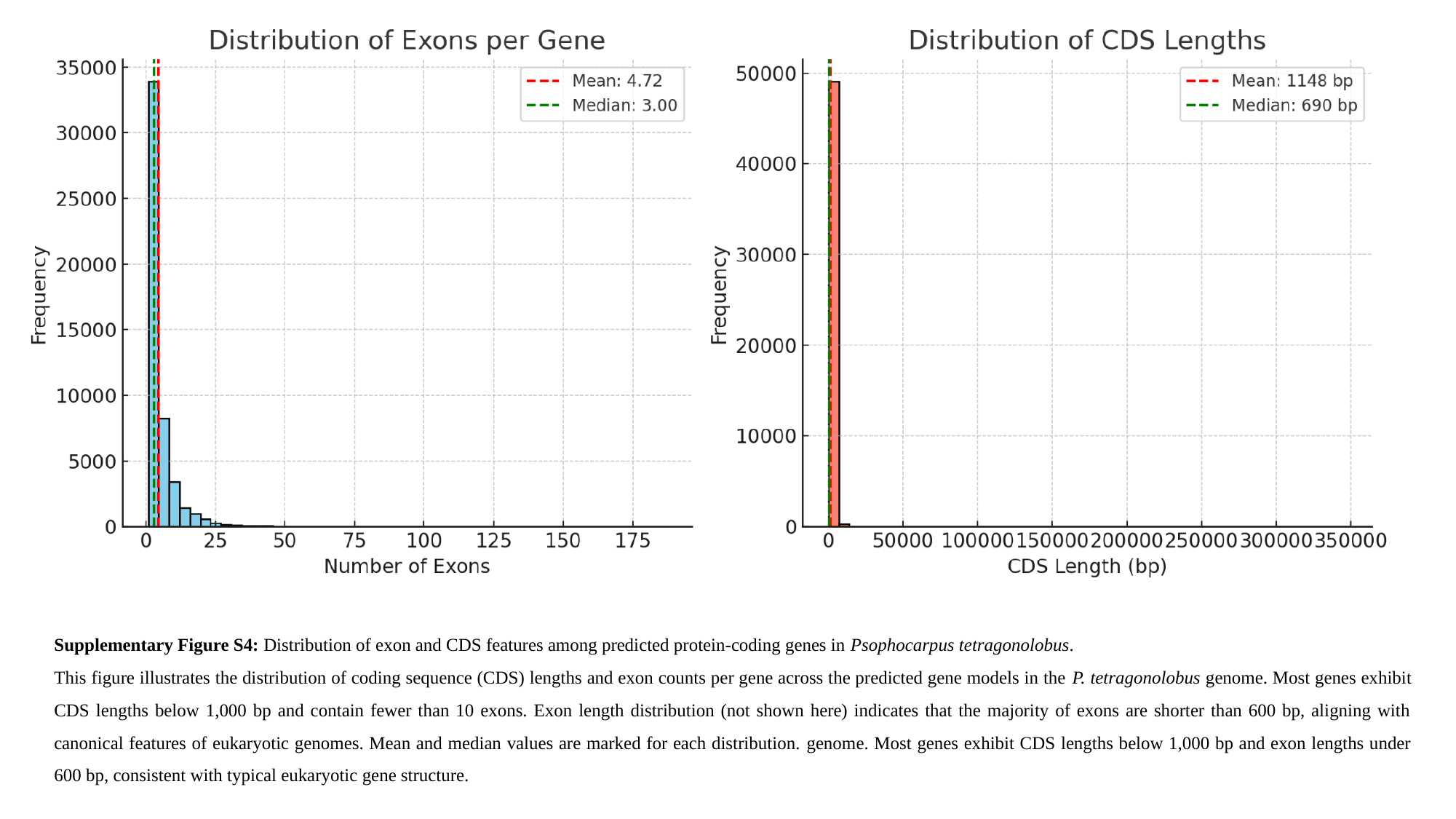

Supplementary Figure S4: Distribution of exon and CDS features among predicted protein-coding genes in Psophocarpus tetragonolobus.
This figure illustrates the distribution of coding sequence (CDS) lengths and exon counts per gene across the predicted gene models in the P. tetragonolobus genome. Most genes exhibit CDS lengths below 1,000 bp and contain fewer than 10 exons. Exon length distribution (not shown here) indicates that the majority of exons are shorter than 600 bp, aligning with canonical features of eukaryotic genomes. Mean and median values are marked for each distribution. genome. Most genes exhibit CDS lengths below 1,000 bp and exon lengths under 600 bp, consistent with typical eukaryotic gene structure.
