## Supplementary figure S5 for "Chromosome-Scale, Telomere-to-Telomere Assembly of Winged Bean (*Psophocarpus tetragonolobus* (L.) DC.) Genome Reveals Lipid Biosynthetic Hotspots and Gene Family Dynamics"

### Slide 1
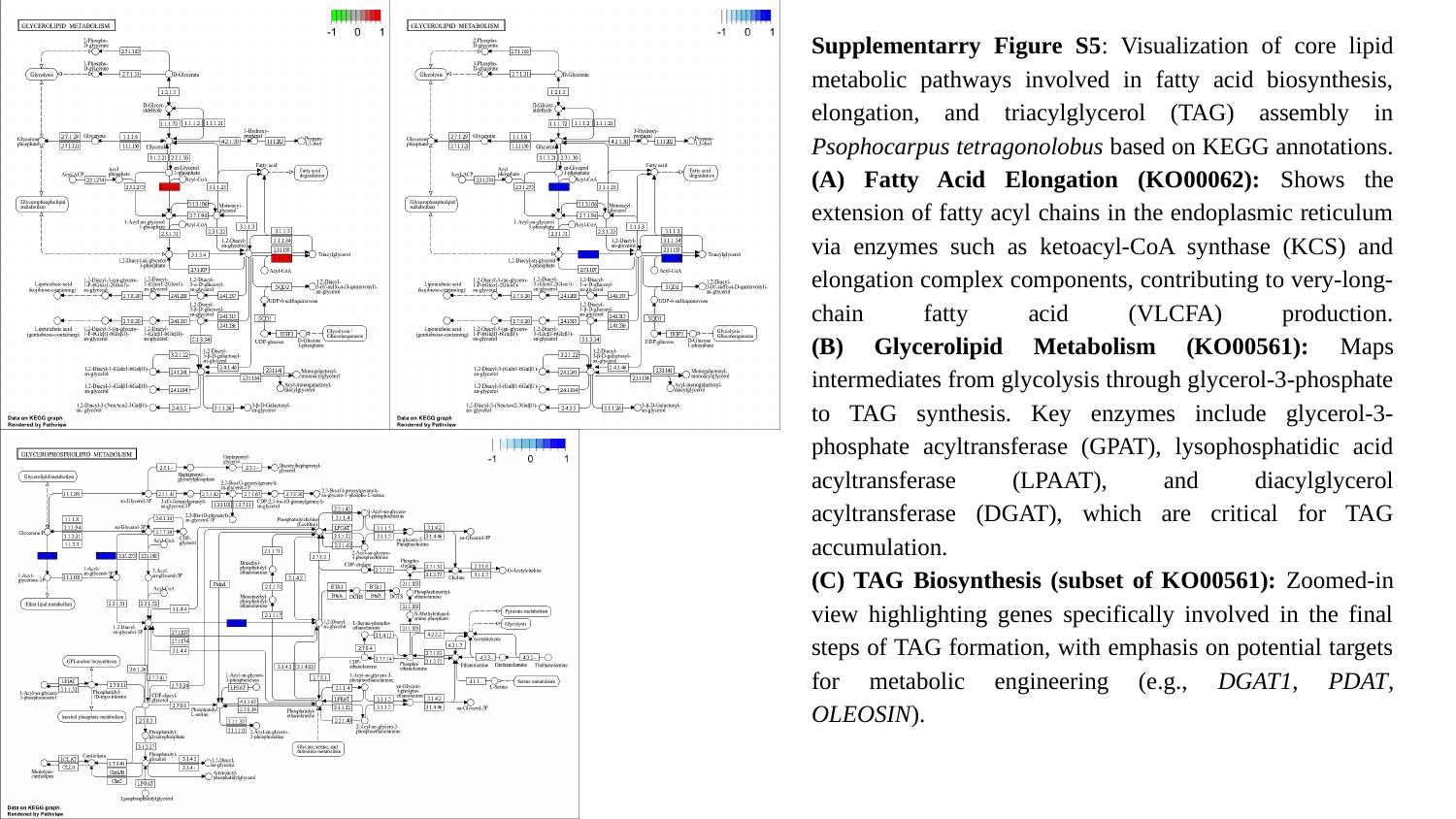

Supplementarry Figure S5: Visualization of core lipid metabolic pathways involved in fatty acid biosynthesis, elongation, and triacylglycerol (TAG) assembly in Psophocarpus tetragonolobus based on KEGG annotations.
(A) Fatty Acid Elongation (KO00062): Shows the extension of fatty acyl chains in the endoplasmic reticulum via enzymes such as ketoacyl-CoA synthase (KCS) and elongation complex components, contributing to very-long-chain fatty acid (VLCFA) production.
(B) Glycerolipid Metabolism (KO00561): Maps intermediates from glycolysis through glycerol-3-phosphate to TAG synthesis. Key enzymes include glycerol-3-phosphate acyltransferase (GPAT), lysophosphatidic acid acyltransferase (LPAAT), and diacylglycerol acyltransferase (DGAT), which are critical for TAG accumulation.
(C) TAG Biosynthesis (subset of KO00561): Zoomed-in view highlighting genes specifically involved in the final steps of TAG formation, with emphasis on potential targets for metabolic engineering (e.g., DGAT1, PDAT, OLEOSIN).
