## Supplementary figure S6 for "Chromosome-Scale, Telomere-to-Telomere Assembly of Winged Bean (*Psophocarpus tetragonolobus* (L.) DC.) Genome Reveals Lipid Biosynthetic Hotspots and Gene Family Dynamics"

### Slide 1
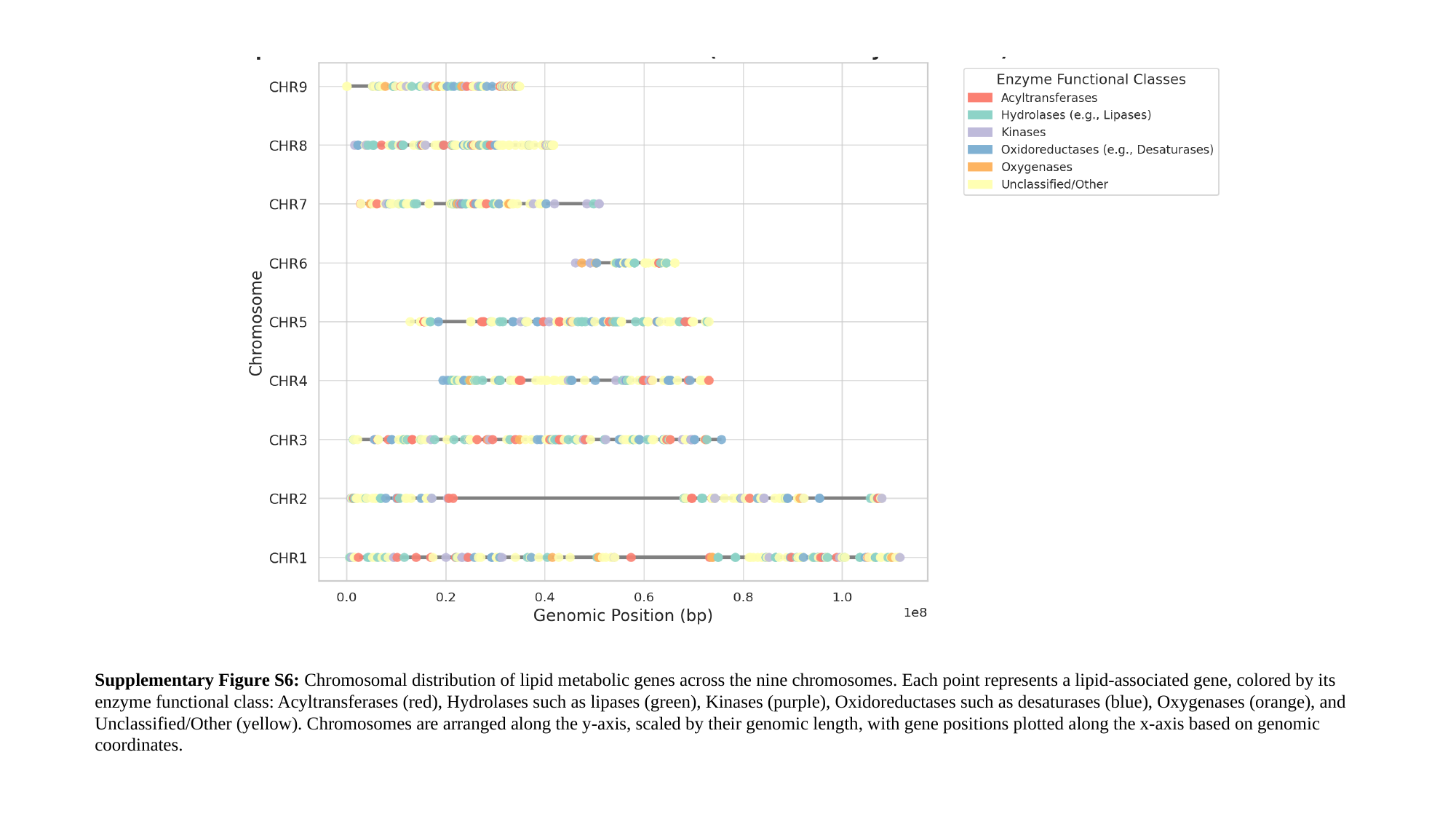

Supplementary Figure S6: Chromosomal distribution of lipid metabolic genes across the nine chromosomes. Each point represents a lipid-associated gene, colored by its enzyme functional class: Acyltransferases (red), Hydrolases such as lipases (green), Kinases (purple), Oxidoreductases such as desaturases (blue), Oxygenases (orange), and Unclassified/Other (yellow). Chromosomes are arranged along the y-axis, scaled by their genomic length, with gene positions plotted along the x-axis based on genomic coordinates.
