## Supplementary figure S7 for "Chromosome-Scale, Telomere-to-Telomere Assembly of Winged Bean (*Psophocarpus tetragonolobus* (L.) DC.) Genome Reveals Lipid Biosynthetic Hotspots and Gene Family Dynamics"

### Slide 1
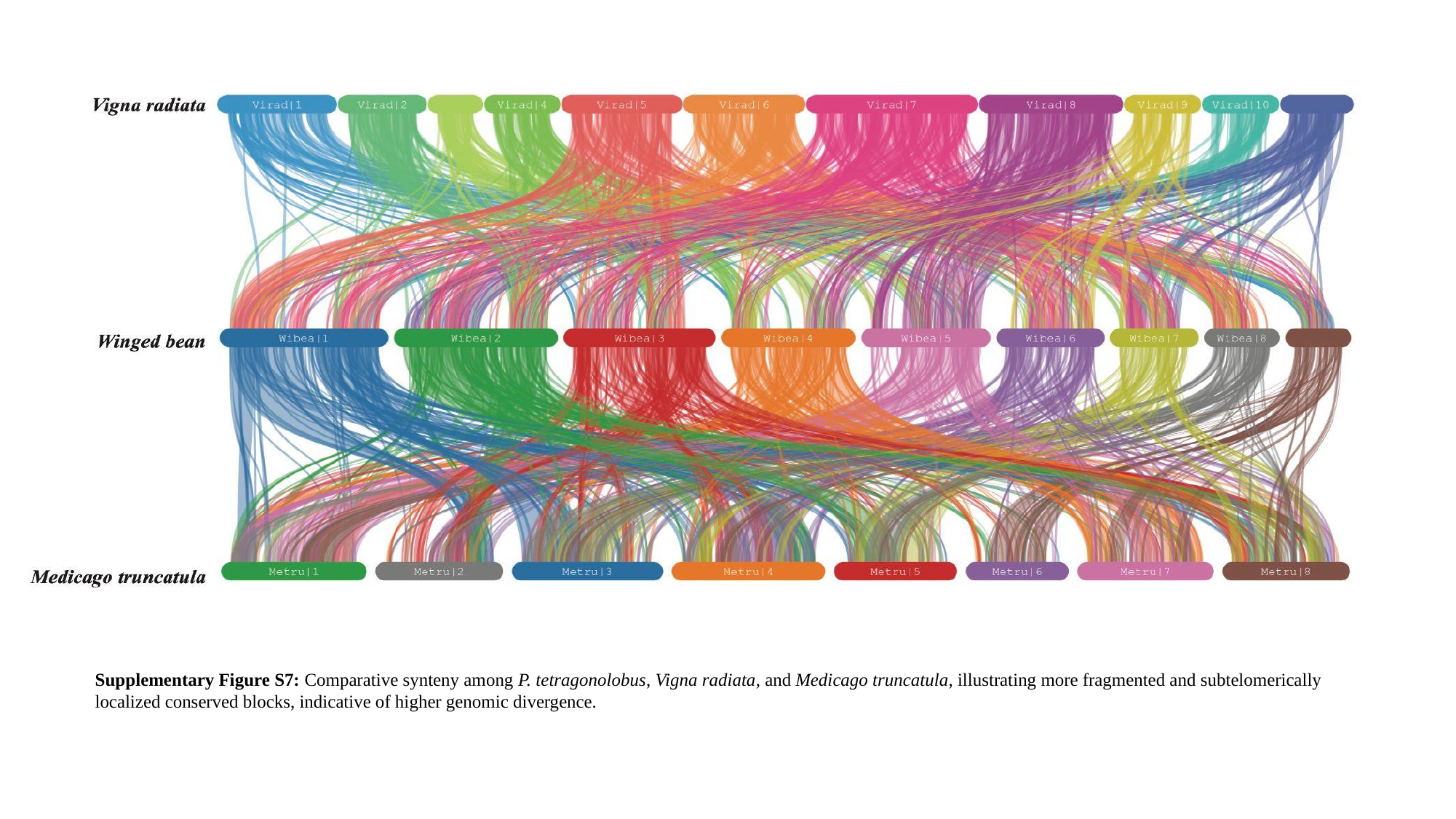

Supplementary Figure S7: Comparative synteny among P. tetragonolobus, Vigna radiata, and Medicago truncatula, illustrating more fragmented and subtelomerically localized conserved blocks, indicative of higher genomic divergence.
